# AlphaFold2 *knows* some protein folding principles

**DOI:** 10.1101/2024.08.25.609581

**Authors:** Liwei Chang, Alberto Perez

## Abstract

AlphaFold2 (AF2) has revolutionized protein structure prediction. However, a common confusion lies in equating the *protein structure prediction* problem with the *protein folding problem*. The former provides a static structure, while the latter explains the dynamic folding pathway to that structure. We challenge the current *status quo* and advocate that AF2 has indeed learned some protein folding principles, despite being designed for structure prediction. AF2’s high-dimensional parameters encode an imperfect biophysical scoring function. Typically, AF2 uses multiple sequence alignments (MSAs) to guide the search within a narrow region of its learned surface. In our study, we operate AF2 without MSAs or initial templates, forcing it to sample its entire energy landscape — more akin to an *ab initio* approach. Among over 7,000 proteins, a fraction fold using sequence alone, highlighting the smoothness of AF2’s learned energy surface. Additionally, by combining recycling and iterative predictions, we discover multiple AF2 intermediate structures in good agreement with known experimental data. AF2 appears to follow a “local first, global later” folding mechanism. For designed proteins with more optimized local interactions, AF2’s energy landscape is too smooth to detect intermediates even when it should. Our current work sheds new light on what AF2 has learned and opens exciting possibilities to advance our understanding of protein folding and for experimental discovery of folding intermediates.

---

AlphaFold2 (AF2) revolutionized the protein structure prediction field (*1*). Its success has led to an array of applications, and assessment of the transferability to other problems such as protein-protein and protein-peptide prediction (*2, 3, 4*). Some efforts have focused on assessing the limits and understanding what AF2 has learned to increase its versatility and applicability. For instance, modifying multiple sequence alignments (MSAs) leverages different co-evolutionary signals in AF2 to identify multiple biologically relevant states (*5, 6, 7*). AF2 was prevalent in CASP15 (Critical Assessment of Structure Prediction) for single protein structure prediction, where differences in prediction accuracy were primarily due to the quality of MSAs used within AF2 (*8*). AF2’s performance has lead many to question whether the protein folding problem has been solved (*9, 10, 11*).

While the protein structure prediction problem focuses on generating static images of a protein’s folded state, the protein folding problem seeks to understand the dynamic processes involved in folding, including the pathways and intermediates that occur. Experimentally characterizing these folding pathways and intermediates is challenging due to the short transition times and the need for high-resolution data. Although computational methods based on physical and chemical principles offer a theoretical approach, they are limited by the accuracy of force fields and the extensive timescales required to simulate folding trajectories (as shown in Fig.1, right). In this study, we explore the capability of AF2 beyond its traditional role in structure prediction, investigating its potential to provide insights into protein folding intermediates.

**Figure 1:**
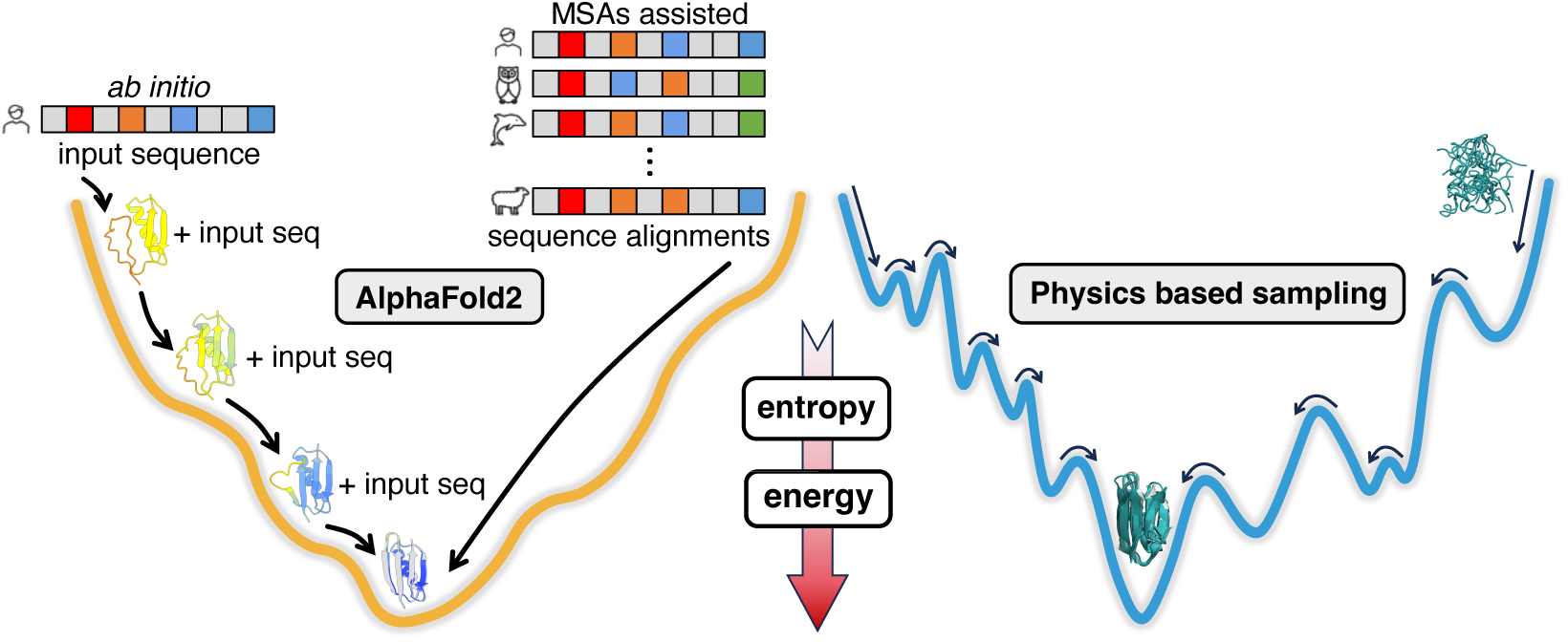
Cartoon representation of the folding energy landscape comparing AlphaFold2’s smooth surface (left) with a rugged landscape typical of physics-based force fields (right). In physics-based approaches, extensive sampling is required to identify the native basin, whereas in AF2, MSAs or protein templates quickly guide structure refinement to narrower regions of conformational space, effectively bypassing other regions. We employ an AF2-*ab initio* approach, iteratively generating structures with the help of last round prediction to navigate the smooth energy function learned by AF2 starting from single sequence alone.

A recent study proposes that AF2 has learned an approximate biophysical energy function for structure prediction, where the co-evolutionary signal from multiple sequence alignments (MSAs) is necessary to find a native conformation with low energy, and the structural module further provides refined structure predictions (*12*). By back-propagating the structural loss and using gradient descent optimization to perturb the input sequence, the model can improve structure prediction accuracy without MSAs. This suggests that AF2 can act as an “energy minimizer” to iteratively improve the quality of the structure prediction. Previously, we observed that AF2 can detect the nuance of local interaction effect in alternating protein folding pathways for a set of four closely related proteins, in agreement with ϕ and ψ – value analysis data and extensive molecular dynamics (MD) simulations (*13*). Here, we propose that the predicted conformations along iterative structure predictions with AF2 could be representative of protein folding intermediates.

We first select a handful of proteins whose folding processes have been widely characterized by experiments and computer simulations – protein G, protein L, ubiquitin, and SH3. The original protein sequence is used to construct the sequence representation followed by an iterative structure prediction process. In the initial step of the iteration, neither template structure information nor MSAs are used. In subsequent iterations, the sequence information is combined with the last predicted structure as a template for next round prediction. As a further application of the methodology, we apply this protocol on the mutants of protein G and protein L to identify whether AF2 could detect changes in folding routes between the wild-type and mutant sequences. We then apply this protocol to designed sequences for these folds, showing that indeed they have smoother folding routes with less frustrations in AF2 compared to their original sequence. Finally, we assess the transferability of this approach by going beyond these six proteins and querying a large set of proteins representative of different folds and sizes from the Protein Data Bank (PDB) (*14*).

## Results

### MSAs or templates can facilitate *sampling* AF2’s learned energy landscape

While MSAs remain a good way to navigate AF2’s learned energy surface, it is not necessarily the only way. For instance, testing a set of decoy structures into the model without co-evolutionary information showed discriminating behavior for structure quality (*12*). This ability to forego MSAs and use structural data has also been applied to enhance prediction accuracy using single-sequence queries with a generator-discriminator approach. They first generate an AF2 structure prediction for the query sequence in the generator model, and predicted structure serves as the input of a discriminator model. The loss of discriminator model is then used by the generator model to update query sequence with gradient descent for the next round of prediction. In this way structures of higher quality from single sequence alone are generated.

### Iteratively *sampling* AF2’s energy landscape slows down structure prediction, but can yield important insights into AF2’s folding funnel

Inspired by their findings, we sought to evaluate the potential of AF2 for predicting folding intermediates through iteratively using query sequence and structure prediction. Here, AF2 predicts a structure from our single sequence query, and the prediction will be combined together with the sequence query for the next iteration (see Fig. 1, left). There are two routes in AF2 to achieve this. One is its internal recycling scheme which is a key design to gradually increase structure prediction accuracy (*1*). Additionally, similar to how the template structure is processed here (*12*), we can use the prediction from each round as distograms and combine it with the pair representation from primary sequence to perform another prediction, which we call an iteration. Recycling and iteration steps can be used either independently or simultaneously. The prediction is repeated until the convergence of both structure prediction and its confidence scores. Thus, this approach foregoes the use of MSAs and focuses on examining AF2’s ability to navigate its learned energy surface from single sequence to the native state.

### AF2 correctly predicts protein folding intermediates for six small proteins

The structures generated along iterative prediction of AF2 for six small proteins (protein G, L and their mutants, ubiquitin, and the SH3 domain) are in excellent agreement with experimentally known intermediates and even transition states (see detailed description in Supplementary Text). Notably, the average pLDDT scores, which indicate prediction confidence, increased as the structures approached their native conformations. Another common trend across these proteins is that early intermediates with native-like regions exhibited low pLDDT scores, which increased as other structural elements fall into place even though their conformations remain the same (see Fig. 2 and 3). Proteins G and L share a common topology despite of low sequence similarity and different folding pathways. Their mutants have different kinetics and the mutant of protein G even alternates folding pathways of its wild type. Our method revealed specific folding intermediates consistent with previous experimental and computational findings (*15, 16, 17, 13, 18, 19, 20, 21, 22, 23, 24, 25, 26*), demonstrating AF2’s capacity to capture intricate folding pathways. In protein G, the iterative structure predictions show a folding pathway starting with a native C-terminal hairpin formation, followed by a registry-shifted N-terminal hairpin, before proceeding to find the native fold. AF2 is sensitive to sequence changes introduced in protein G_*mut*_, folding through the N-terminal with a more locally favorable turn than the native sequence after specific mutations, in good agreement with experiments. Similarly, protein L starts folding through the N-terminal hairpin first, with subsequent iterations gradually folding the C-terminal hairpin. Once more, AF2 detects a much smoother folding landscape for protein L_*mut*_ that follows a similar pathway as the original sequence. Some subtle but important details about their folding pathways observed previously can be easily identified from our iterative predictions, such as the formation of a registry-shifted N-terminal hairpin in transition state which then refolds to native in protein G (*17*) and the folding of C-terminal hairpin in later step because of the low stability in its turn region for protein L (*20*). Our findings also extend to ubiquitin and the SH3 domain, where AF2 successfully predicted folding intermediates that match known observations (*27, 28, 29, 30, 31, 32, 33, 34*). Ubiquitin’s AF2 folding pathway exhibits early formation of an N-terminal hairpin, and samples different C-terminal conformations before establishing correct end-to-end contacts which lead to the folded state – reflecting previous findings. For the SH3 domain, the iterative method preserved the middle three β-strands early on, with N- and C-terminal contacts forming later.

**Figure 2:**
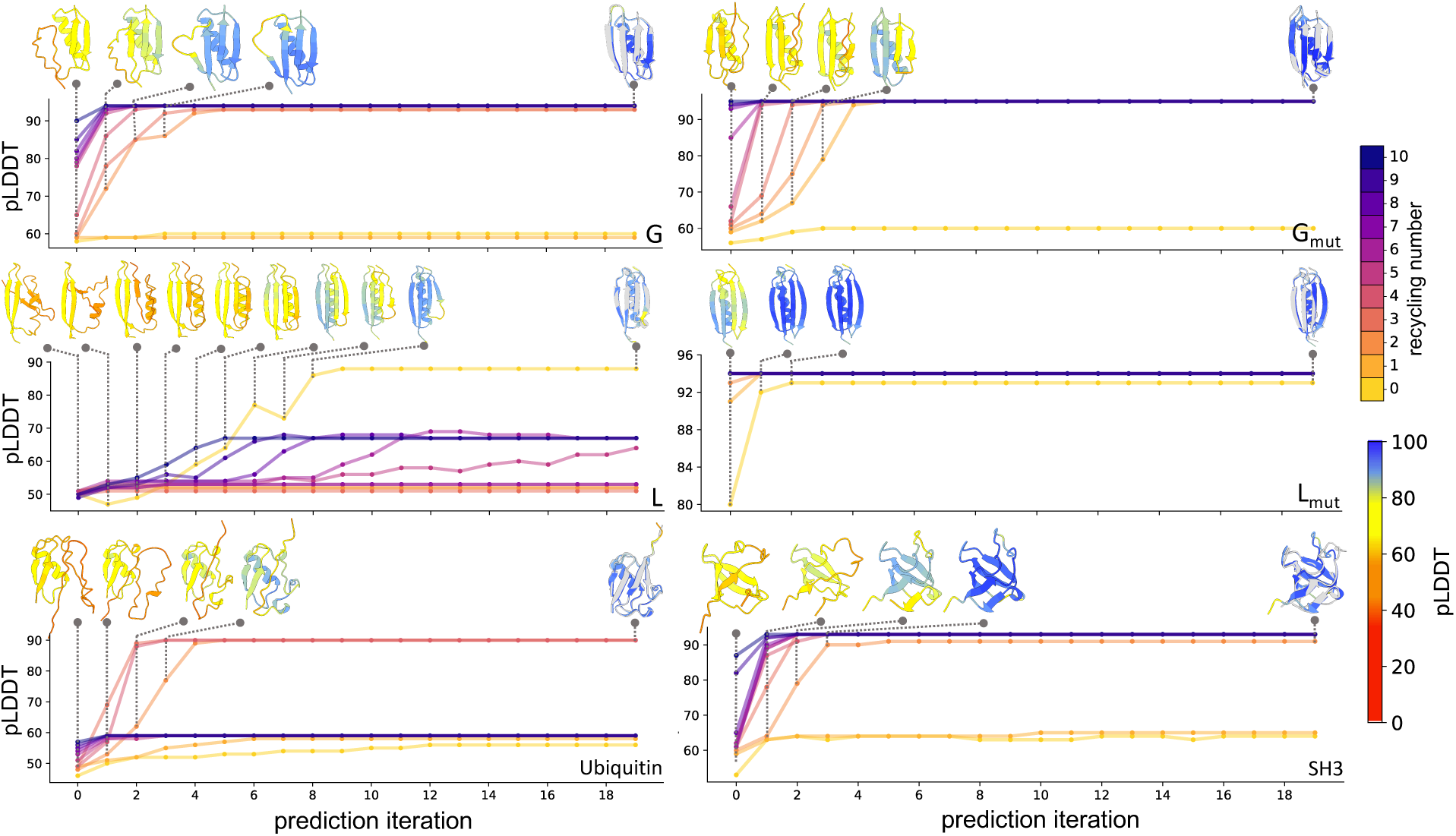
Iterative structure prediction of proteins G, L and their mutants, ubiquitin, and SH3. Each subplot represents predictions for each protein with increasing number of recycling for AF2 from 0 to 10. The structure predictions are shown for the iteration that finally finds the native state with the least recyclings. All structures before converging to the native and the structure at the 20th iteration aligned with the native (colored in grey) in that iteration are depicted (colored by pLDDT scores).

**Figure 3:**
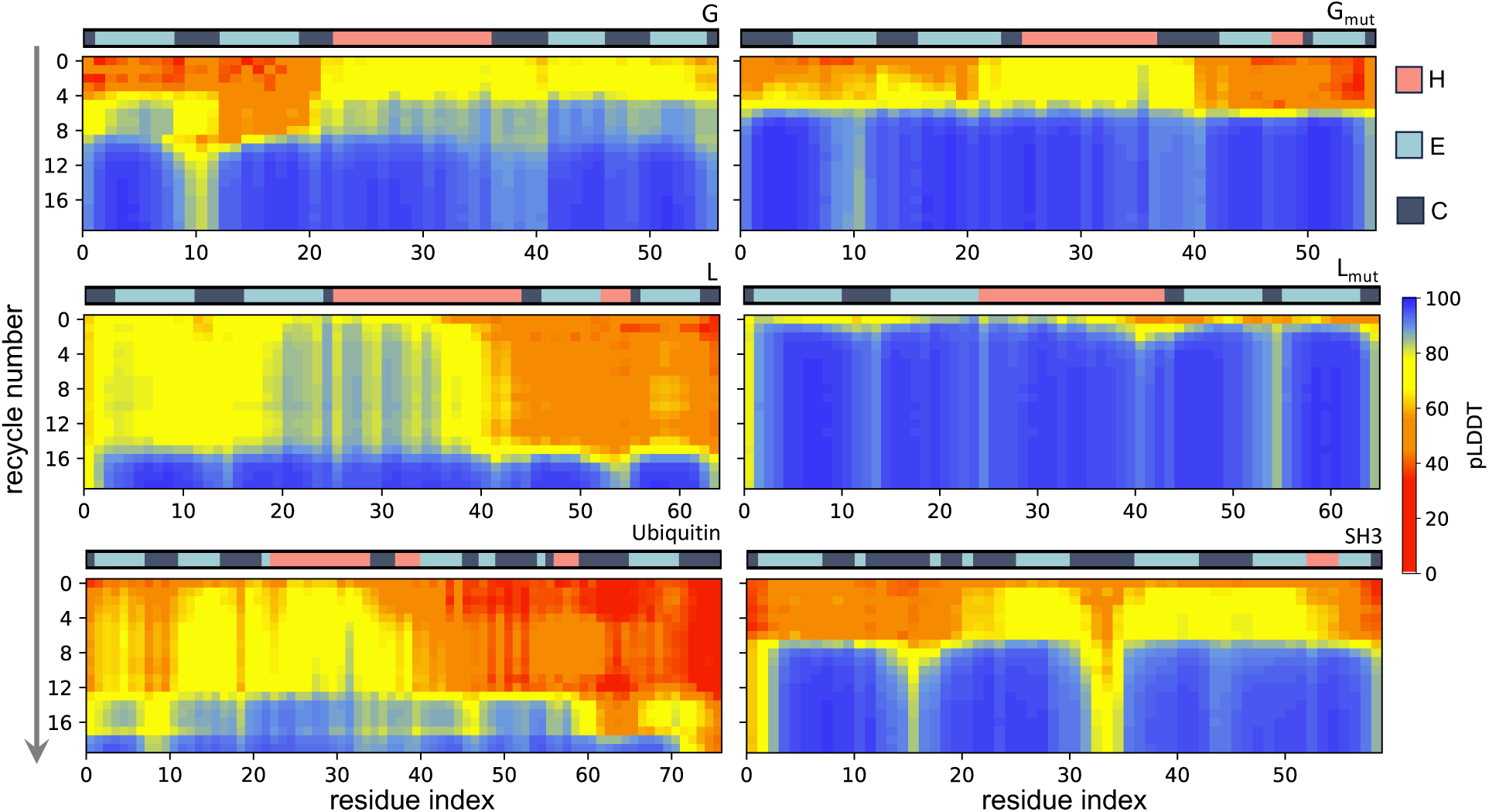
AlphaFold2 single sequence based prediction with recycling only. Each plot represents the evolution of structure prediction in terms of residue wise pLDDT along with the number of recyclings. The secondary structure classification of native structure is depicted above each plot.

Most α-helices in these systems are predicted to fold and unfold either fully or partially through MD simulations and are only stabilized once they pack against other native structural elements. However, AF2 typically predicts these helices early on, even with low pLDDT scores (see Fig. 2 and 3). Such “overstabilization” of α-helix in AF2 was also reported in independent studies (*3*).

### Iterative structure predictions follow a “local first, global later” folding mechanism

One aspect of the protein folding problem explores whether a universal principle governs the folding patterns of most proteins, while also providing specific behavior for each unique sequence.

Levinthal’s paradox suggests the existence of a physical principle that prevents the exploration of all possible conformations because the search process would be impractical timewise (*35*). Folding funnels offer an explanation that folding happens as the free energy decreases while the remaining conformational space for searching is reduced (*36, 37*).

One common pattern from the above iterative structure predictions is that these six proteins tend to follow a “local first, global later” folding mechanism. We measure conformations predicted with this iterative approach by three metrics (see Fig. 4) – average contact order (<*CO*>, the average native contacts formed by the inter-residue distances along the sequence), average effective contact order (<*ECO*>, differs with <*CO*> in that the inter-residue distance considers both spatial and sequence effects), and the ratio of short and long range contacts (see Methods in SI). The initial iterations favor local contacts, as seen by low <*CO*> and <*ECO*> scores. This leads to new contacts that remain close in terms of <*ECO*>, but are higher in <*CO*>. Effectively, once a contact is established, it brings residues that are far in the sequence (high <*CO*>) close in space (low <*ECO*>), reducing the entropic penalty for conformational search process. The ratio of short and long range contacts delivers the same message that structure predictions favor short interactions in the beginning, which facilitates the formation of longer range contacts in later iterations. For smaller size proteins that fold, such as the widely studied fast-folding proteins, the peptide chain tends to follow a collapse-condensate mechanism, which shows the semi-folded structure resembles the final shape that was driven by a hydrophobic collapse and others (*21*). As the length of the protein sequence increases, chances for establishing long-range interactions become smaller. Thus, short-range interactions that are more prevalent in the unfolded state can reduce conformational search space to assist the folding of the overall topology.

**Figure 4:**
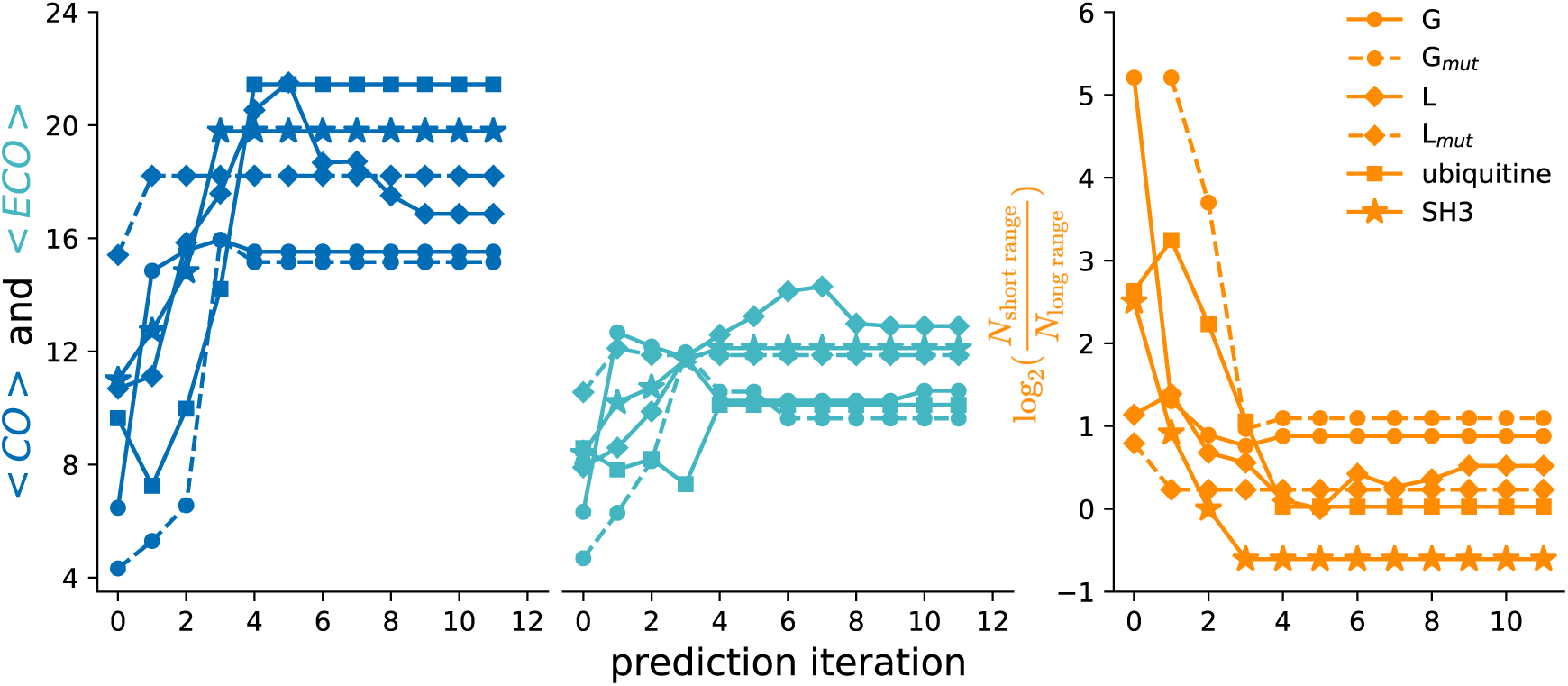
Local and global effect in iterative structure predictions with AlphaFold2 measured by average contact order (<*CO*>, dark blue), average effective contact order (<*ECO*>, blue-green), and the ratio of short and long range contacts (dark orange).

### Generated sequences by ProteinMPNN encode smoother energy landscapes with more optimized local interactions than naturally occurring sequences

Machine learning models trained on existing knowledge of protein structures and sequences have been successful in predicting novel sequences given a desired template topology. Sequences generated from ProteinMPNN, for example, show high correlation between pLDDT from AF2 single sequence prediction and true LDDT-Cα against template structure (*38*). The prevalence of more favorable local interactions in designed sequences over natural ones can be indicative that evolutionary pressure does not need to over-stabilize every local structure element.

To test this hypothesis, we predicted 20 new sequences with ProteinMPNN based on the native structure for the six targets. For each one of them, we repeated the iterative structure prediction strategy. Supplementary figures 2-13 showcase our results for the evolution of pLDDT score and RMSD against the template backbone using AF2 models 1 and 2 during iterative structure predictions. For most of the sequences, AF2 is able to predict structures resembling the template topology after only a few iterations, independent of the number of recycles. The folding funnels are smooth enough that AF2 is able to predict the structures in a single leap, with no detected potential intermediates. Among the six targets, Protein G mutant presents the smoothest iterative predictions, where all 20 sequences rapidly found the conformation they were designed from. SH3 has a larger number of exceptions, where the number of recycles affects the iterative predictions, and we see more failures in predicting native-like structures. Similarly, we also see this happens in a few cases for the remaining proteins. However, future experiments are required to distinguish whether this is because they are failed sequence designs by ProteinMPNN or their native structure cannot be predicted with AF2 using our single sequence based approach.

### Scaling AF2 iterative structure prediction to diverse protein folds

General principles of protein folding have been sought by theories and experiments for decades. Multiple factors were proposed in accurately predicting protein folding rates including contact order, packing compactness, and secondary structure compositions (*39, 40, 41, 42*). However, none of the existing models can provide the folding mechanism of each individual protein at atomistic level. Continuous development of computer simulation methods with ever-increasing computing power led to good agreement with experimental observations, but their capability is mostly limited to small globular proteins. We took one step further to scale our iterative structure prediction with AF2 on the known protein folds. The sequences were chosen from PDB by the following criteria: (1) we downloaded sequences of protein monomer structures deposited by March 5th, 2023 with length ranging from 30 to 250; (2) we filtered sequences whose deposited model has a resolution higher than 3 Å ; (3) we clustered the remaining sequences using the *easy-linclust* tool provided in *MMseqs2* with default options. Overall, we collected 7418 sequences from the PDB to perform iterative structure prediction with AF2. For each sequence, we run iterative predictions with recyclings 0, 1, 3, 5, and 8 for 500 iterations. Fig. 5A shows the first two dimensions of t-SNE embedding for our selected protein space after converting each native structure into a topology based feature vector using Gauss Integral (*43*). This plot shows the diversity of selected protein folds in secondary structures and the ability of iterative structure prediction to correctly fold a small subset of sequences (success indicated by the final structure closer than 3 Å to the native state in the middle right plot of Fig. 5A). We also found that sequences sharing similar folds with SH3 and ubiquitin in PDB are close to each other in the embedding. Not surprisingly, AF2’s ability to fold proteins through an iterative approach is inversely proportional to protein size (Fig. 5B). In particular, for fragments below 50 residues, the success rate is around 20%, which helps explain AF2’s success in predicting the bound structure of peptides without MSAs (*3*). However, for proteins over 100 residues, the success rate rapidly falls below 5%. We run predictions with both models that can take template structures in AF2 - models 1 and 2, which were fine-tuned with different number of extra sequences and training samples. We can see that the best structure prediction of each model after 500 iterations differs in terms of RMSD against their native structure for many targets (Fig. 5C). The secondary structure distribution of our curated subset from PDB shows several trends that can be representative of all protein structures in PDB: (1) the percentage of coil-like fragments that appear in this clustered subset is around 36%, (2) protein structures with all β-sheets are rare, and (3) most structures have α-helix between 0 ∼ 40%, while structures with larger portions of α-helix are also frequent (see Fig. 5D). Naturally, as a trained model with structures in PDB, AF2 tends to predict well for α-helix rich structures but fails for structures with more coil fragments (see Fig. 5E). Such prediction bias towards α-helix in this large-scale prediction is consistent with our observation of their appearance during the early stage along iterative predictions above (Fig. 2,3).

**Figure 5:**
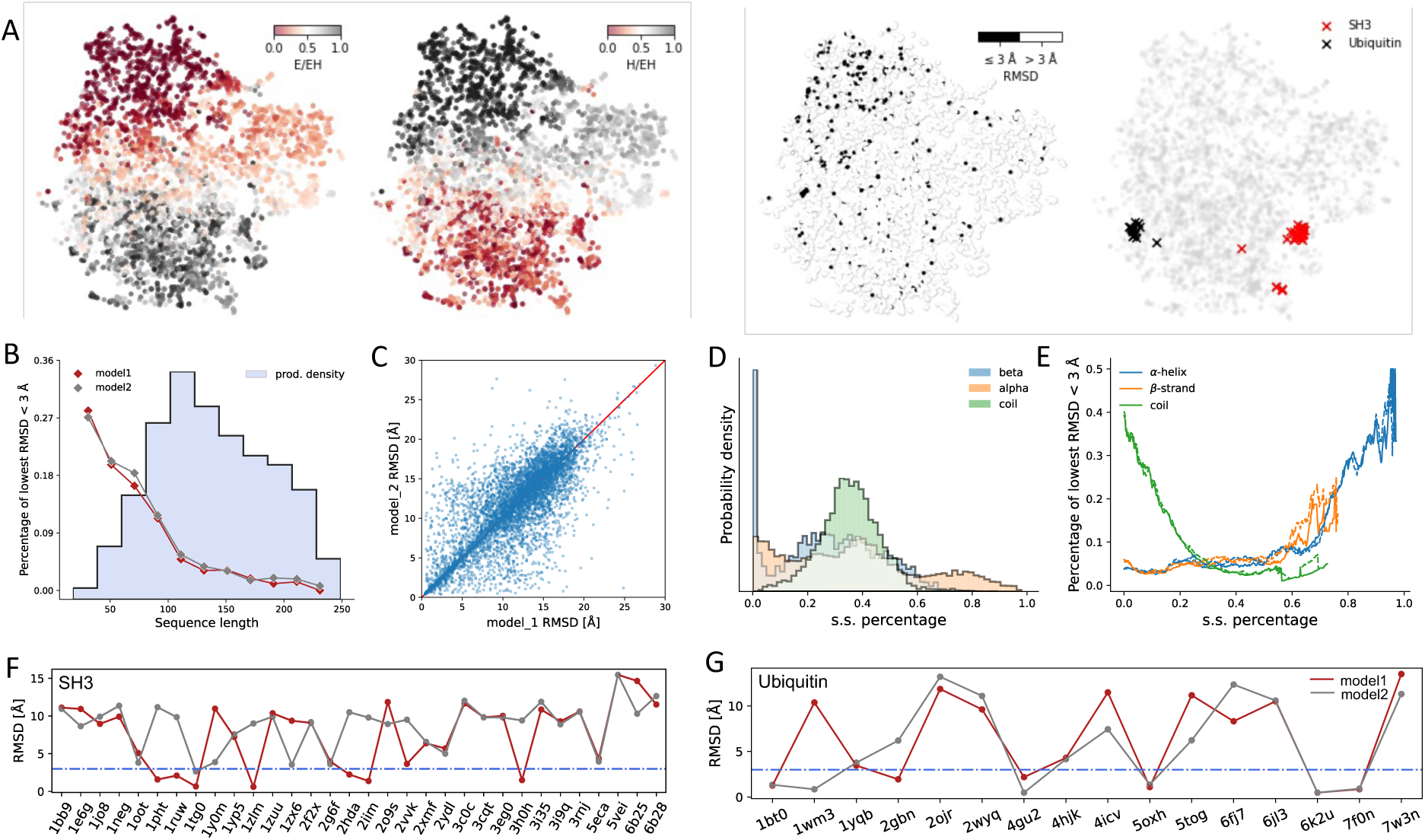
Large scale iterative structure predictions for 7418 proteins curated from the PDB. **A.** structural feature based t-SNE embedding plot colored based on four different properties: ratio of β-sheets (left) and α-helix (middle left) over the sum of both, sequences the AF2-*ab initio* folds to within 3 Å of native (middle right), and positions of SH3 (red) and ubiquitin (black) like proteins (right). **B.** Percentage of proteins folding into structure less than 3 Å from native decreases with longer sequences for both predictions by model 1 (red) and model 2 (grey). The sequence length distribution of all selected proteins is shown in blue histogram. **C.** Comparison of predictions from model 1 and 2 in terms of the lowest RMSD against native structure from all iterative predictions. **D.** The percentage of secondary structure distribution of each type for all selected structures. **E.** Percentage of proteins fold into structure less than 3 Å from native versus the amount of secondary structure for each type. **F & G.** The lowest RMSD from iterative predictions for SH3 (panel F) and ubiquitin (panel G) like proteins by model 1 (red) and 2 (grey).

### Discovery of folding intermediates in SH3, ubiquitin-like proteins, and beyond

An intriguing aspect of the protein folding problem is determining whether proteins with the same fold share similar folding mechanisms. This can reveal whether differences in sequence lead to alternative folding pathways that converge on the same final conformation. In our study of proteins G, L, and their mutants, we observed that subtle sequence changes in protein G could enhance local interactions, altering the folding pathways from which the native conformation is achieved. In contrast, the folding pathway for protein L and its mutant remained consistent despite sequence variations. With the large-scale iterative predictions, we extended our investigation to other proteins with different folds. We discovered that structures with folds similar to SH3 and ubiquitin cluster closely together in structure-based embeddings (see Fig. S14 and 5A, right). However, despite sharing the same conformation overall (see their structural alignment in Fig. S14), not all proteins within each fold type could be accurately predicted, and the two AF2 models produced differing results (see Fig. 5F, G). Among all SH3 and ubiquitin like proteins, only a few achieved conformations with less than 3 Å from the native structure. Despite low sequence similarity, these seven proteins appear to follow the same folding pathway as the SH3 protein we studied above (PDB: 2HDA), where the middle three β-sheets stack together first, awaiting the formation of interactions between the N- and C-terminal strands (Fig. S15). Predictions from ubiquitin like proteins also demonstrate that they likely have similar folding intermediates where the first two β-strands tend to fold early on, followed by the assembly of native interactions at both termini (Fig. S16). However, we are unsure whether those SH3 and ubiquitin-like proteins that did not find their native structure after iterative predictions fold in different pathways or also share similar folding patterns.

### AF2 identifies differential stability in Fibronectin type III domain repeats

Fibronectin (FN) type III, a key component in extracellular matrices, is a 368 residue protein whose modular domain topology is foldable in AF2 through our current single sequence approach. FNIII is composed of four modules called repeats, numbered 7 to 10 (^III^7-10). Our iterative structure predictions with recycling number greater or equal to one obtain final structures in close agreement with the experimental structure (*44*), with a small deviation in dihedrals between ^III^7 and ^III^8 (see Fig. 6A). Interestingly, AF2 finds significant differences in the folding of the different domains. With domain ^III^7 finding the native structure rapidly (single iteration), while ^III^8 and ^III^10 remain partially folded, and ^III^9 remains unfolded. In subsequent steps, both ^III^8 and ^III^10 fold, and so does most of the ^III^9 repeat – but the overall assembly of the domains is incorrect, and AF2’s confidence score remains low. After a few more iterations, the linkers between ^III^8-9 and ^III^9-10 are accurately predicted, but the one between ^III^7-8 (linker 1) keeps changing the orientation during later iterations without entering the native conformation. We hypothesize this is because the linker 1 region is more flexible than the other two linkers. Although detailed descriptions of the folding pathways for this four-repeat protein are not available, several studies have reported similar observations.

**Figure 6:**
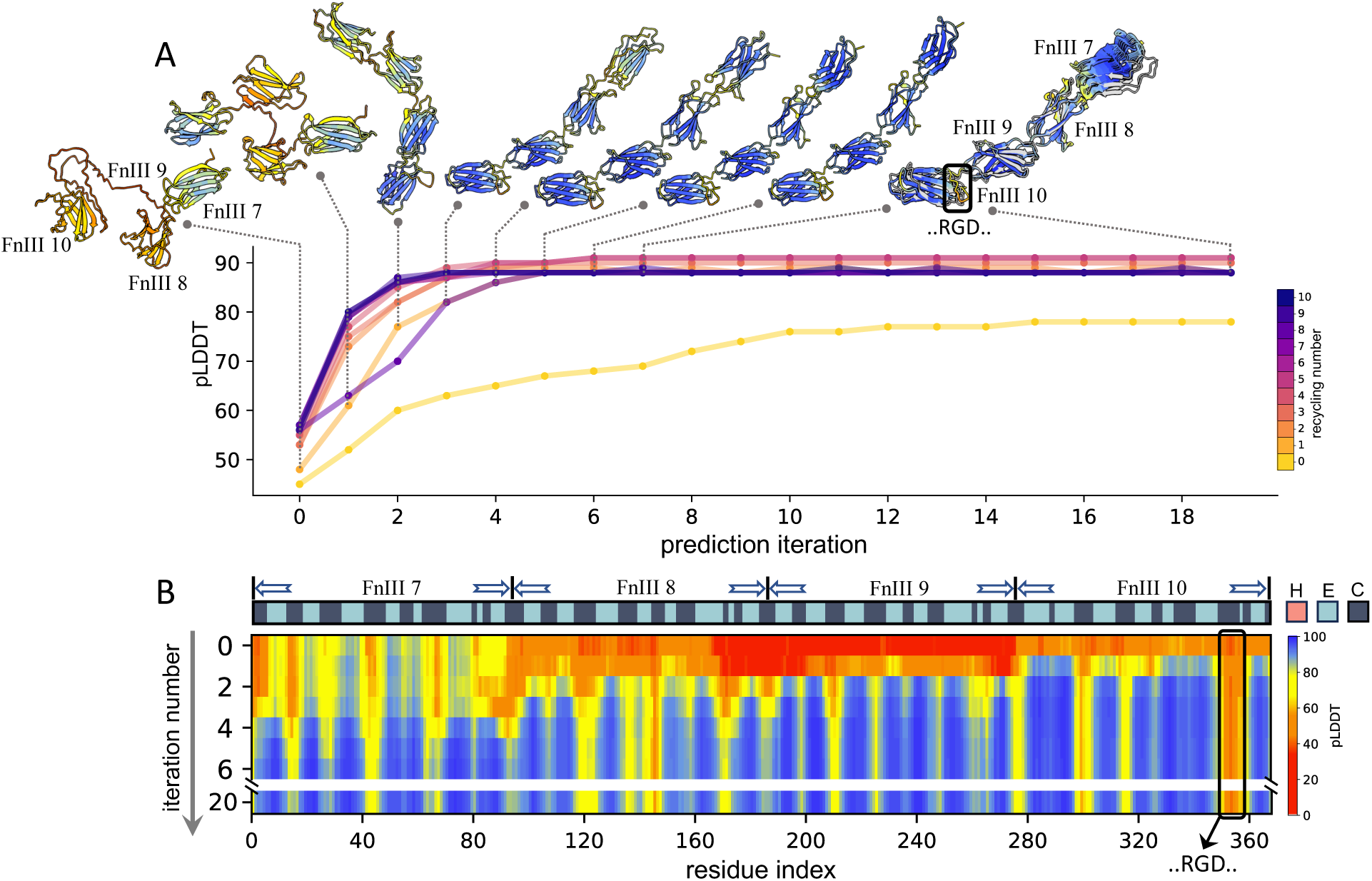
A. Iterative structure predictions of Fibronectin type III with AlphaFold2 using different number of recycles. The repeat names and location of integrin binding loop containing RGD peptides are labeled. B. The evolution of structure prediction in terms of residue wise pLDDT along representative iterations. The secondary structure classification of native structure is depicted above.

Experimental studies from (*45,46*) indicate that ^III^9 is the least stable one among the four repeats with equilibrium stability (Δ*G* _*f*_ ) of -1.2(±0.5) kcal mol^−1^ compared to -6.1(±0.1) kcal mol^−1^ for ^III^10, and the loop region containing the sequences Arg–Gly–Asp (RGD) from the ^III^10 can bind integrin themselves. This aligns with our iterative predictions in that ^III^9 is unstructured in the beginning and the last repeat to find native conformation while the RGD region remains flexible with low pLDDT scores throughout the iterative predictions. It has also been suggested that ^III^9 and ^III^10 act independently in that the mutations of one domain have no effect on the other (*47*). Additionally, a thorough experimental study on the dynamics and stability of Fibronectin recently found that ^III^7 appears to have no effect on the stability of the rest of modules since ^III^7-10 and ^III^8-10 are comparably stable while ^III^8 was found to help stabilize ^III^9 and ^III^10 (*48*). Those experimental evidence help us explain why AF2 is able to come up with a conformation resembling the crystal structure despite of its long sequence length overall: (1) the folding of all four repeats are mostly driven by local interactions in each repeat, similarly for the linkers in between, and (2) the more stable repeat likely has more locally favorable interactions so ^III^7 and ^III^10 can be predicted early on while the least stable ^III^9 folded at the end. This indicates that our iterative structure prediction can help provide the relative stability information among the Fibronectin repeats.

## Discussion

Unraveling the protein folding process at an atomic level is critical for understanding protein functions and their roles in diseases. This task largely depends on two intertwined factors: a highly accurate energy function that describes intricate molecular interactions and efficient sampling methods that can quickly explore the vast conformational space of proteins, which have high degrees of freedom. Traditional computational simulations often struggle with inaccuracies in force fields and insufficient conformational sampling. Even when extensive sampling is achieved, distilling insights about protein folding pathways can be challenging, as some methods may bias the folding route (e.g., through restraints in MELD (*49*) or the use of fragments in Rosetta (*50*)). Recovering statistically significant pathways is complex, as shown in our extensive studies of protein G, protein L, and their mutants using various MD simulation methods and others (*13, 51, 52*).

Recent advancements have shown that the millions of trained parameters in AF2 can act as a sort of biophysical energy function that AF2 navigates to predict structure (*12*). Our study aims to enhance our understanding of what AF2 has learned about protein structures and explore whether this learned energy surface can be leveraged to simulate the protein folding process. Unlike traditional physics-based methods (e.g., sampling along atomistic or coarse-grained force fields), AF2’s energy function appears to be smoother and more precise when native-like interactions are found. It has been particularly successful in using multiple sequence alignments (MSAs) to identify regions of phase space for structure determination (*1, 5, 6, 7*), likely due to co-evolutionary information providing shortcuts to different conformational spaces.

However, our results indicate that in the absence of MSAs, AF2 still exhibits predictive power using only sequence information, resembling more of an *ab initio* approach. Not surprisingly, the structure prediction success rate decreases with increasing sequence length, a limitation that contrasts with the success of *ab initio* methods like Rosetta, MELD, or UNRES (*53*), which have been effective for sequences of similar lengths. Interestingly, AF2-*ab initio* is also successful for proteins composed of independently foldable domains despite their longer length with sequence alone, as the case of Fibronectin shows.

The folding pathways of proteins in AF2 seem to follow a “local-first, global-later” approach. AF2 appears to first fold residue pairs that are nearby in sequence without MSAs, and then locks in structural elements as they get closer in space. When MSAs are unavailable, AF2 likely uses structural information from previous iterations to reduce conformational search space. However, this search becomes increasingly difficult as sequence length grows and more disordered regions are introduced. Additionally, AF2’s pLDDT scores, which reflect the confidence in predicted structures, improve as nearby residues become more native-like, suggesting that the per-residue pLDDT score reflects the quality of a fragment within the context of the entire protein.

From a technical point, we are using AF2 in a way that it was not designed to work (no MSAs), for a purpose that it was not intended (pathways and folding intermediates). It is unclear what is the right balance between iterations and recycling and why it works on some protein sequences but not others. Using recycling alone is effective for the six small proteins, but it is slightly less efficient compared to using both together in some cases (Fig. 3). Of over 7,000 proteins we tested, AF2 has higher success chances on the smaller proteins, and those with more secondary structure. However, we can see successful predictions distributed across the embedding that represent diversity in terms of secondary structure and other properties (see Fig. 5). This is interesting in the sense that in the process of learning to predict protein structures, AF2 has learned something deeper about proteins that allows it to fold some small proteins in ways that are compatible with known findings. It is conceivable that future versions of such approaches will learn more and more about the protein folding principles.

In the broader context, evolution does not know about folding pathways or structures; it just has some selection requirements for actions to happen at particular time intervals with robustness and precision. Evolution does not optimize; it just needs things to be good enough. Optimizing interactions might lead to two proteins never unbinding, which might continuously express a gene or limit the release of oxygen into the blood. However, we as a field are learning the principles to optimize sequences that fold robustly and remain stable with tools such as ProteinMPNN. Several designed proteins were introduced in previous CASP events, typically performed well for physics-based *ab initio* methods. Part of it can be explained by optimized local interactions. Not surprisingly, such protein sequences perform extremely well for iterative prediction using AF2 without MSAs, folding with few or no apparent intermediates in the majority of cases.

The transient nature of intermediate states poses a challenge to their experimental characterization and limits our ability to quantify the predicted accuracy of AF2. Even when those intermediate states are well characterized, there is no single metric that can be used to demonstrate the prediction accuracy of folding process. Furthermore, for systems where intermediates are reported, the results are only qualitative, without atomic details. Hence, the lack of data and objective functions increase the challenge of building a machine learning model for predicting protein folding. Currently, AF2- *ab initio* could be a valuable tool for generating hypotheses about intermediates and guiding the placement of probes to experimentally verify or refute those hypotheses.

## Acknowledgments

We are thankful for the use of HiperGator computational resources at the University of Florida.

## Author contributions

L.C. and A.P. designed the research and wrote the manuscript. L.C. performed all computations and analysis.

## Data and materials availability

We provide publicly available scripts at https://github.com/PDNALab/AlphaFolding for running iterative structure predictions with AlphaFold2 within ColabDesign (https://github.com/sokrypton/ColabDesign). A Colab notebook version of the script is currently available at https://github.com/ccccclw/ColabDesign/blob/main/af/examples/alphafolding.ipynb.

## Supplementary Materials

### Methods

#### Average native contact order, average native effective contact order and the ratio of short and long range contacts

A contact is defined by a residue pair *r*_*ij*_ whose pairwise distance *d*_*ij*_ < 6.5 Å. The set of residue pair distances *D* : {*d*_*ij*_} are selected from native structure given this criterion and we calculate their pairwise distances for both native structure *D*_*native*_ and each structure prediction *D*_*prediction*_.

Then average native contact order <*CO*> is defined as 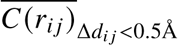 to calculate the average contact order of each contact in predicted structure where its distance is close to the native with the difference Δ*d*_*ij*_ less than 0.5 Å . Similarly, the effective contact order <*ECO*>is defined with a simplified version of (*54*) where each effective contact order is the difference between their contact order and the largest contact order among all native contacts in between. The ratio of short and long range contacts is approximated by the ratio between the number of native contacts being less than 8 residues and more than 16 residues away. In addition, to balance that different secondary structures have varying adjacent residue distance, we weight each residue by their secondary structure type based on average adjacent residue distance for as the following: 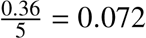 for α-helix type since one helical turn has 5 residues with end-to-end distance roughly 0.36 Å ; similarly, <u>^0.7^</u> = 0.23 for β-strand type and we simply take the average of them for coil type.

### Supplementary Text

#### Detailed description of iterative structure predictions for six small proteins using AF2

We selected six widely studied proteins (protein G, protein L and their mutants, ubiquitin, and the SH3 domain) to assess whether the conformations predicted along this iterative approach resemble folding intermediates found in previous studies. We performed iterative predictions for each protein and compared the effect of changing the amount of recycling done at each iteration. We found that (1) the pLDDT score that measures AF2 prediction confidence increases as the predicted structure progressively enters the native conformation basin; (2) the conformations predicted along the iterations resemble folding intermediates reported in previous works.

Proteins G and L share the same structural topology ( four β-strands forming a β-sheet packed against a helix), despite sharing only about 15% sequence similarity. Despite their structural similarity, their folding pathways are significantly different and have been the object of multiple experimental and computational studies. We and others have shown that protein G has an intermediate state containing the C-terminal hairpin and an additional register-shifted N-terminal hairpin in the transition state due to the local preference of a type I’ turn “NG” over the native type I turn (*13, 15, 16, 17*). Effectively, this register shifted N-terminal hairpin allows contacts between strands one and four, whereby the second strand unfolds and then refolds to native N-terminal hairpin.

Our results using AF2 show that by applying sequence and predicted structure as template information iteratively, the structure of protein G is gradually optimized, as seen by a monotonic increase in global pLDDT score through different iterations (see Fig. 2). Moreover, the conformations in the steps before the native conformation align well with the structural characterization from previous findings of folding intermediates. When the recycling is set to two, the initial prediction from single sequence contains only a structured C-terminal hairpin. Upon feeding this predicted conformation as a template for the next round together with query sequence, the N-terminal hairpin partially emerges in its hairpin-shifted form with the more locally favorable “NG” turn, which later converts to the native turn conformation. By comparison, the protein G mutant (NuG2) contains designed mutations to favor the formation of the native N-terminal turn. Previous investigation from experiments and computer simulations works suggest that such mutations lead to alternative folding routes (*15, 16, 17, 13*). Indeed, we found the iterative prediction using AF2 can capture this behavior clearly, where it first folds the N-terminal hairpin and, in subsequent iterations, folds the C-terminal hairpin.

For protein L, both ϕ and ψ analysis support the formation of an N-terminal hairpin in the transition state, and the appearance of non-native C-terminal hairpin was only found in ψ analysis (*18, 19*). The designed mutation of protein L made the turn region of the C-terminal hairpin more locally stable in the native form, which leads to faster folding speed but doesn’t alternate folding pathways (*20*). Such behavior is again predicted by the iterative prediction from AF2: first, the intermediate with only an N-terminal hairpin is formed, with the formation of the C-terminal hairpin requiring several rounds of iteration for protein L while the mutation in its mutant makes discovery of native state much smoother. Whereas increasing the number of recycles leads faster to native-like structures for protein G and protein G_mut_, this is not the case for protein L. We observe that combining iterations with higher number of recycles leads to stabilizing intermediates that are not further refined (see the middle row of Fig. 2).

The topology of ubiquitin is a combination of five β-strands linked by an α-helix. As in the case of protein G and L, the conclusion from transition state ensemble experimental measurements using ϕ and ψ differs in the appearance of β3 and β4 in addition to the first two strands at N-termini (*27, 28, 29, 30*). An atomic-level description of its folding pathways was provided by MD simulations (*21*) in the millisecond timescale. Among all simulated transition paths, folding always starts with the native form of the N-terminal hairpin, which results in a populated intermediate in an unfolded state, followed by the formation of structural components at the C-termini. The transition state ensemble was found to have both N-termini and C-termini formed in a native-like conformation. Our iterative prediction resembles the description of important states found in both MD simulation and experiments where the first round prediction only contains native contacts at the N-termini with other parts remaining disordered and finding their native conformation after the C-termini *samples* the native interactions (bottom left of Fig. 2). The iterative predictions of ubiquitin also do not follow a “more recycling – faster folding” scheme. In this case, iterations with recycling number set as 2,3,4 entered the native state.

The SH3 domain shares the common structural feature that all five proteins considered above – the N, C-terminal β sheets are close in contact and have five β-strands pairing together in an anti- paralleled way. Experimental studies using ϕ and double-mutant analysis of the folding transition state for SH3 suggests that the β-strands in the middle remain structured and both N and C-terminals stay unstructured (*32, 33*). We observed similar patterns during the iterative prediction with AF2. The iterative predictions exhibit the middle three β-strands conserved from the first step while waiting for the formation of native contacts at the N,C-termini (bottom right of Fig. 2). Akin to the on-pathway and low-populated intermediate structure proposed from previous NMR experiments, the C-termini is the last fragment folded into native state as seen from the pLDDT scores (*34*).

A common trend across these proteins was that early intermediates with native-like regions exhibited low pLDDT scores, which increased as other structural elements fall into place even though their conformations remain the same (see Fig. 2 and 3). Another conclusion from the analysis is that iterations and recycling effectively sample the energy landscape differently. Hence, some runs (e.g., protein L and ubiquitin) using more recycling might lead to locally optimized structures in a minima of AF2’s energy surface from which it cannot recover to sample other topologies. Additionally, we found that all the six proteins can be predicted with only recycling (see Fig. 3 and S1). There is no trend that we could find (high or low number of recycles) that were generalizable. Thus, independent iterative runs at different recycling are recommended. Fortunately, AF2 is always aware of the quality of its predictions based on the pLDDT score – remaining at low pLDDT scores even for part of the structure it cannot further refine.

## Supplementary Figures

**Figure 7: Supplementary figure 1.**
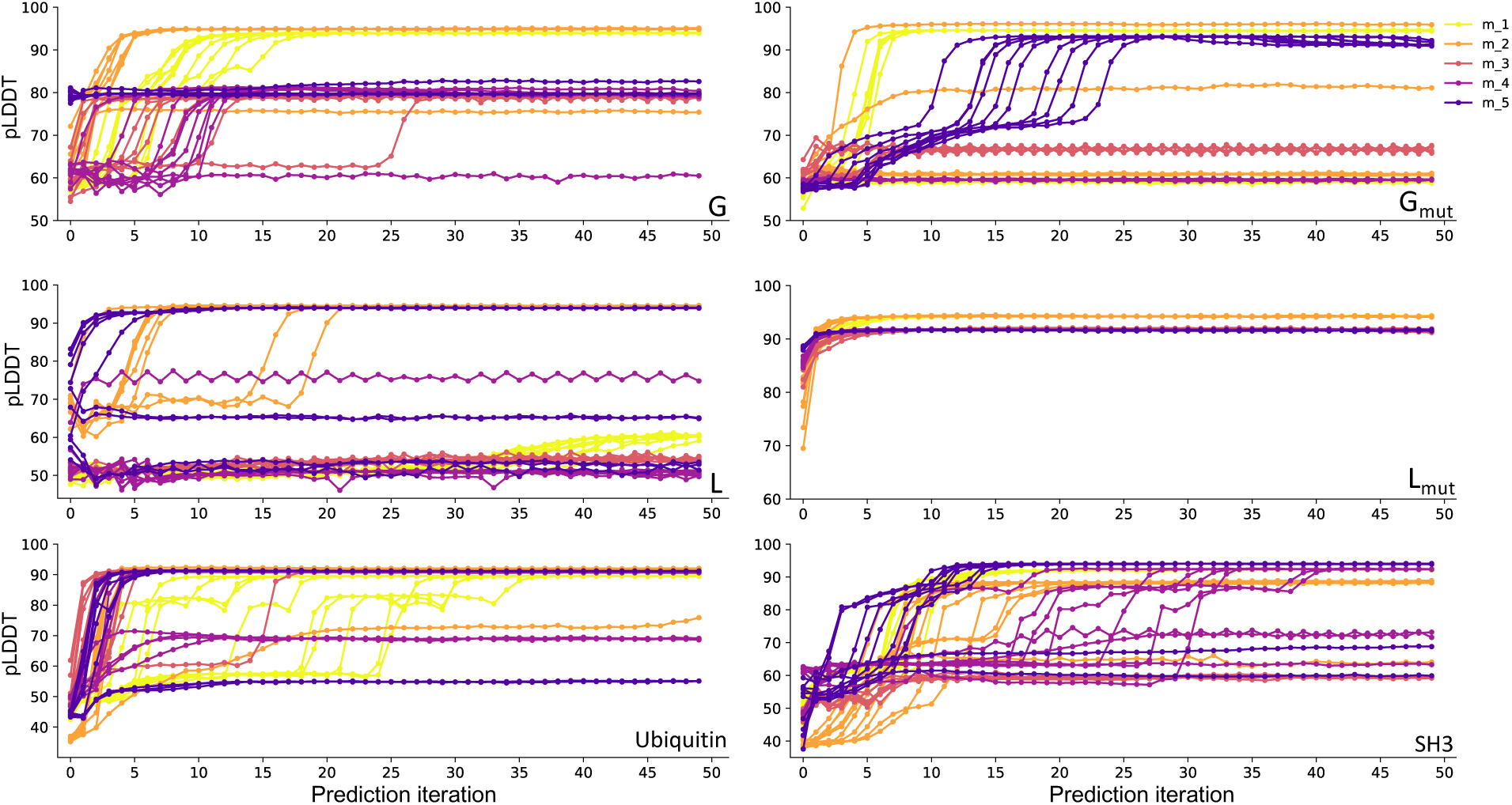
AlphaFold2 single sequence based prediction with recycling only. Each plot represents the evolution of structure prediction in terms of their average pLDDT along with the number of recyclings. We run predictions with different random seeds for all five models in AlphaFold2.

**Figure 8: Supplementary figure 2.**
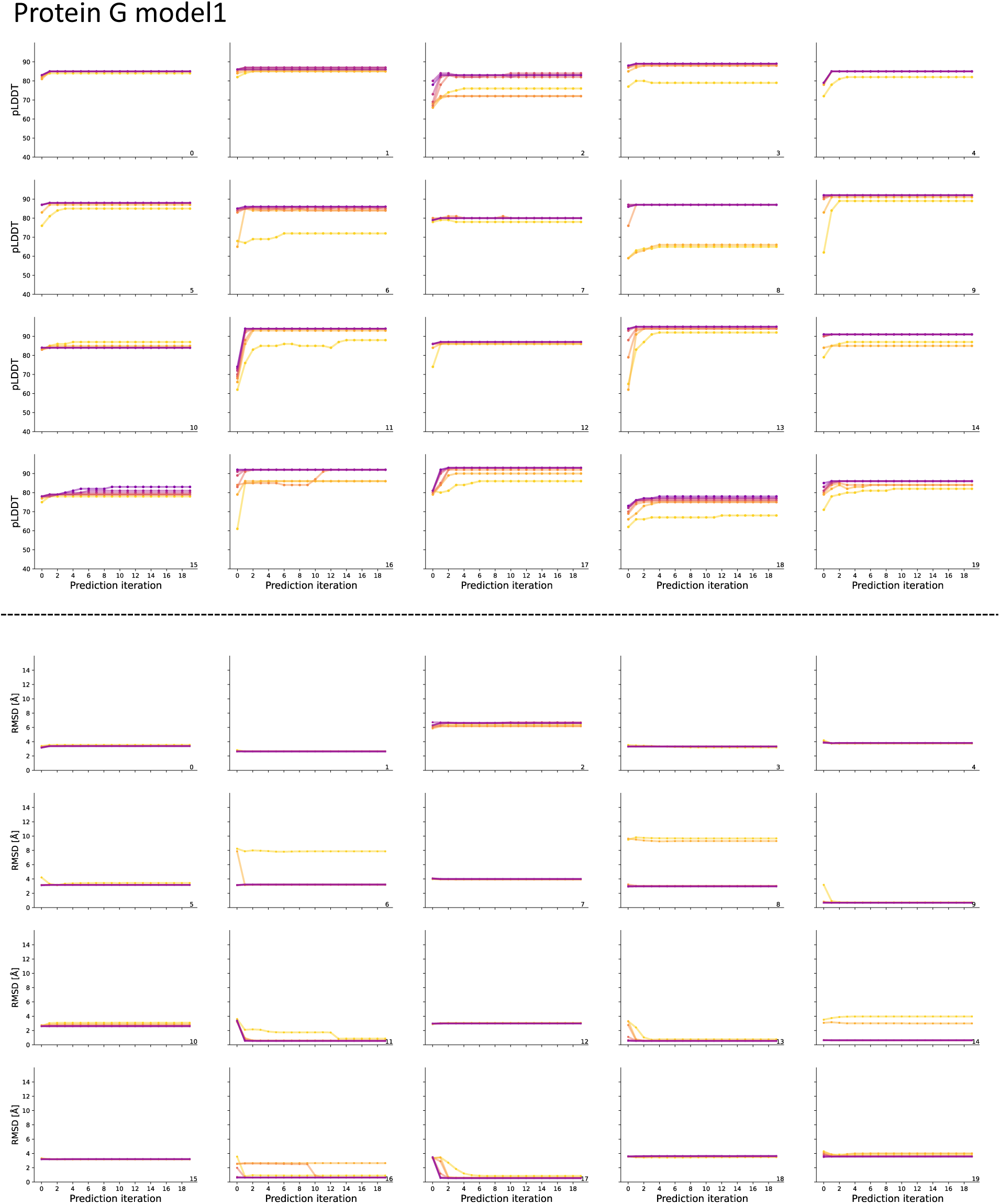
Iterative predictions for 20 ProteinMPNN designed sequences of protein G measured by pLDDT (top) and RMSD (bottom) from native using model 1 of AF2.

**Figure 9: Supplementary figure 3.**
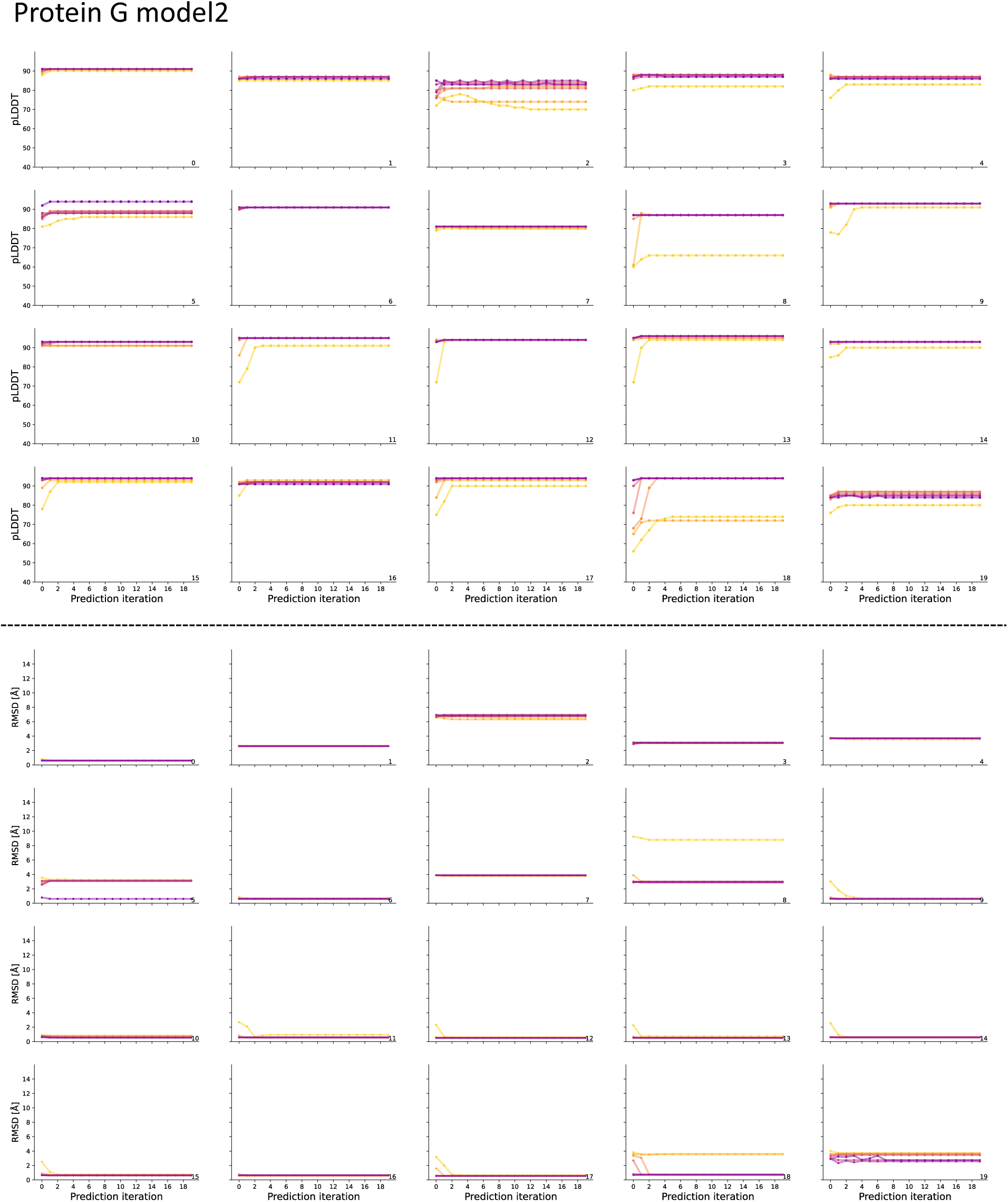
Iterative predictions for 20 ProteinMPNN designed sequences of protein G measured by pLDDT (top) and RMSD (bottom) from native using model 2 of AF2.

**Figure 10: Supplementary figure 4.**
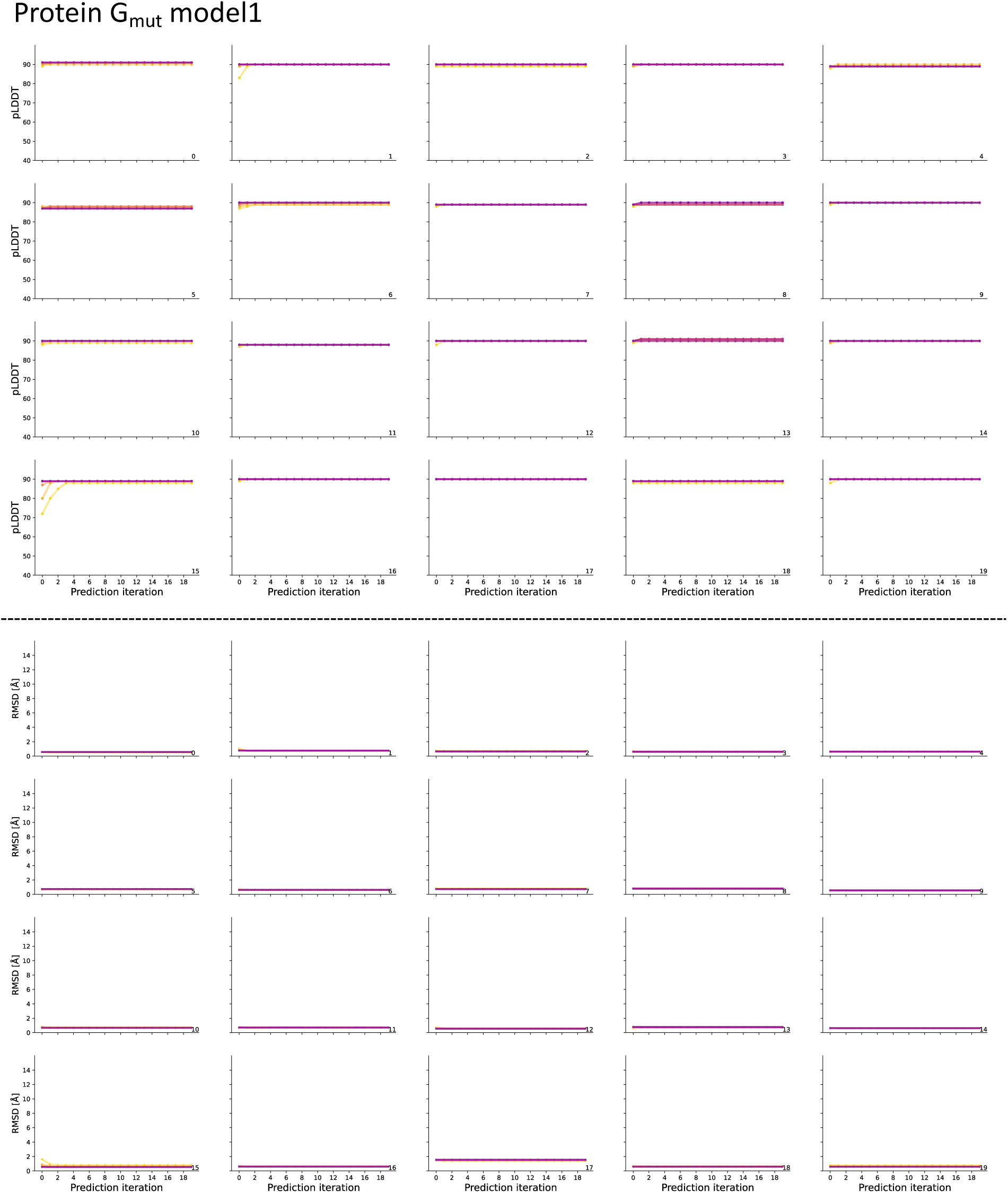
Iterative predictions for 20 ProteinMPNN designed sequences of protein G mutant measured by pLDDT (top) and RMSD (bottom) from native using model 1 of AF2.

**Figure 11: Supplementary figure 5.**
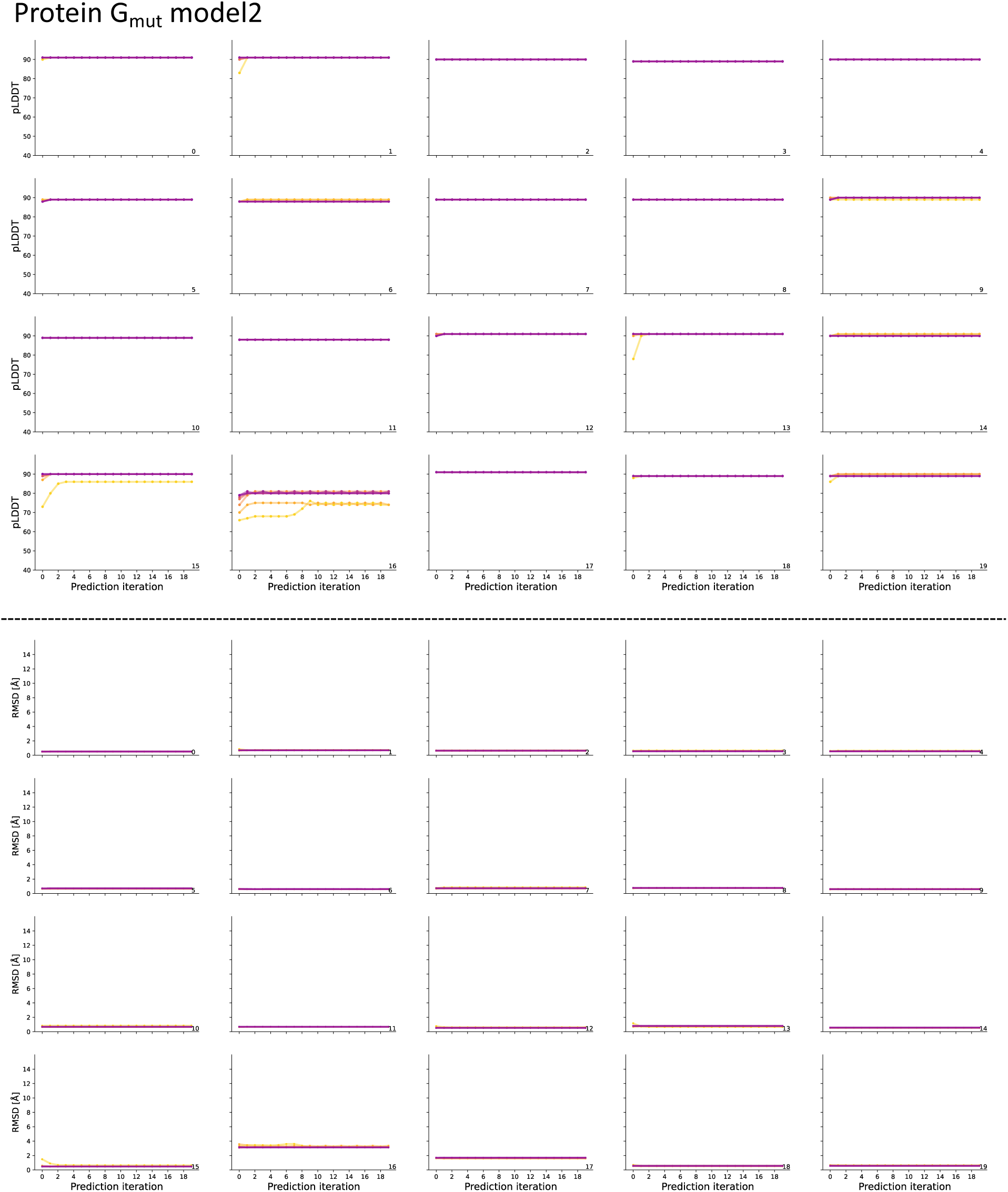
Iterative predictions for 20 ProteinMPNN designed sequences of protein G mutant measured by pLDDT (top) and RMSD (bottom) from native using model 2 of AF2.

**Figure 12: Supplementary figure 6.**
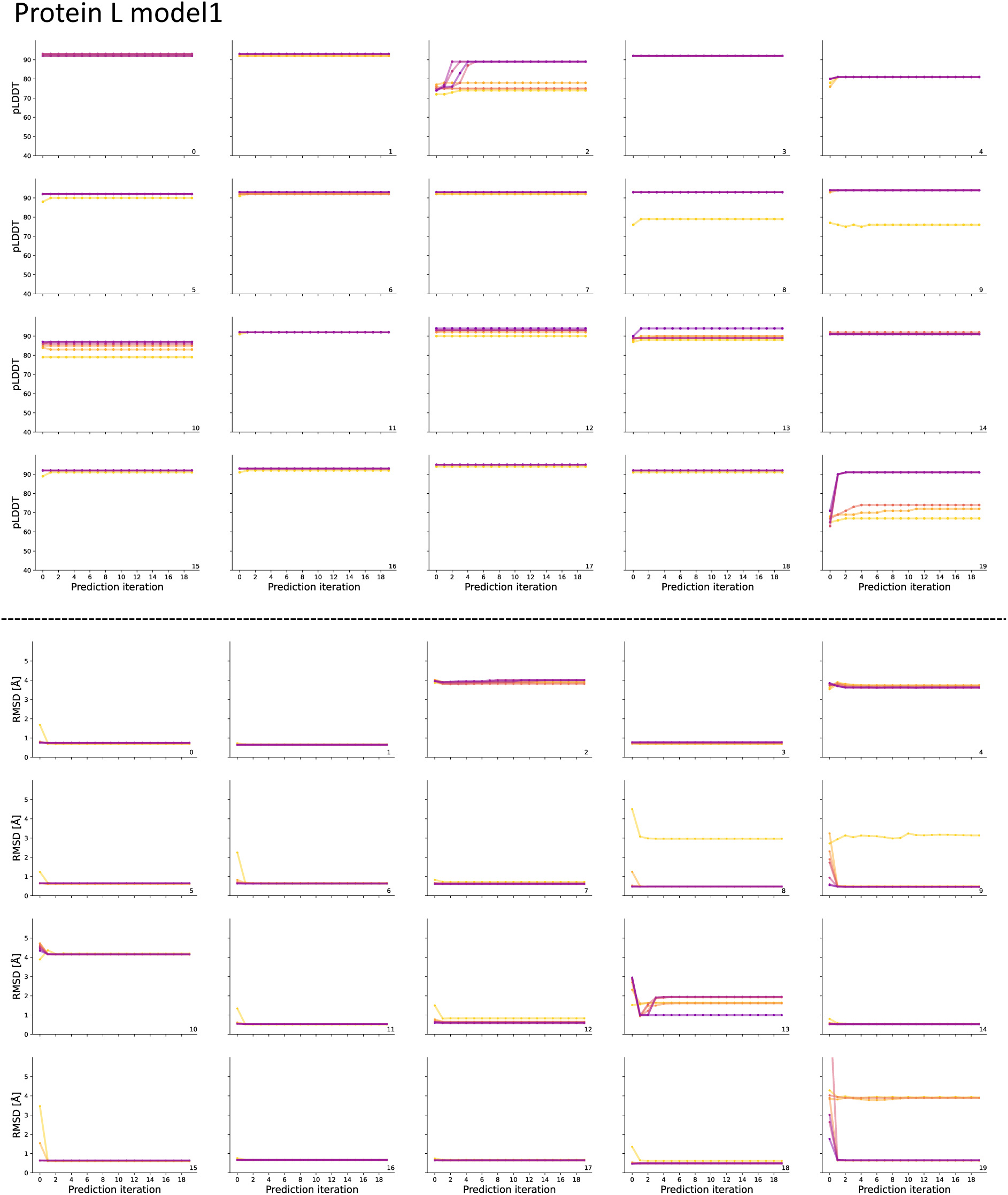
Iterative predictions for 20 ProteinMPNN designed sequences of protein L measured by pLDDT (top) and RMSD (bottom) from native using model 1 of AF2.

**Figure 13: Supplementary figure 7.**
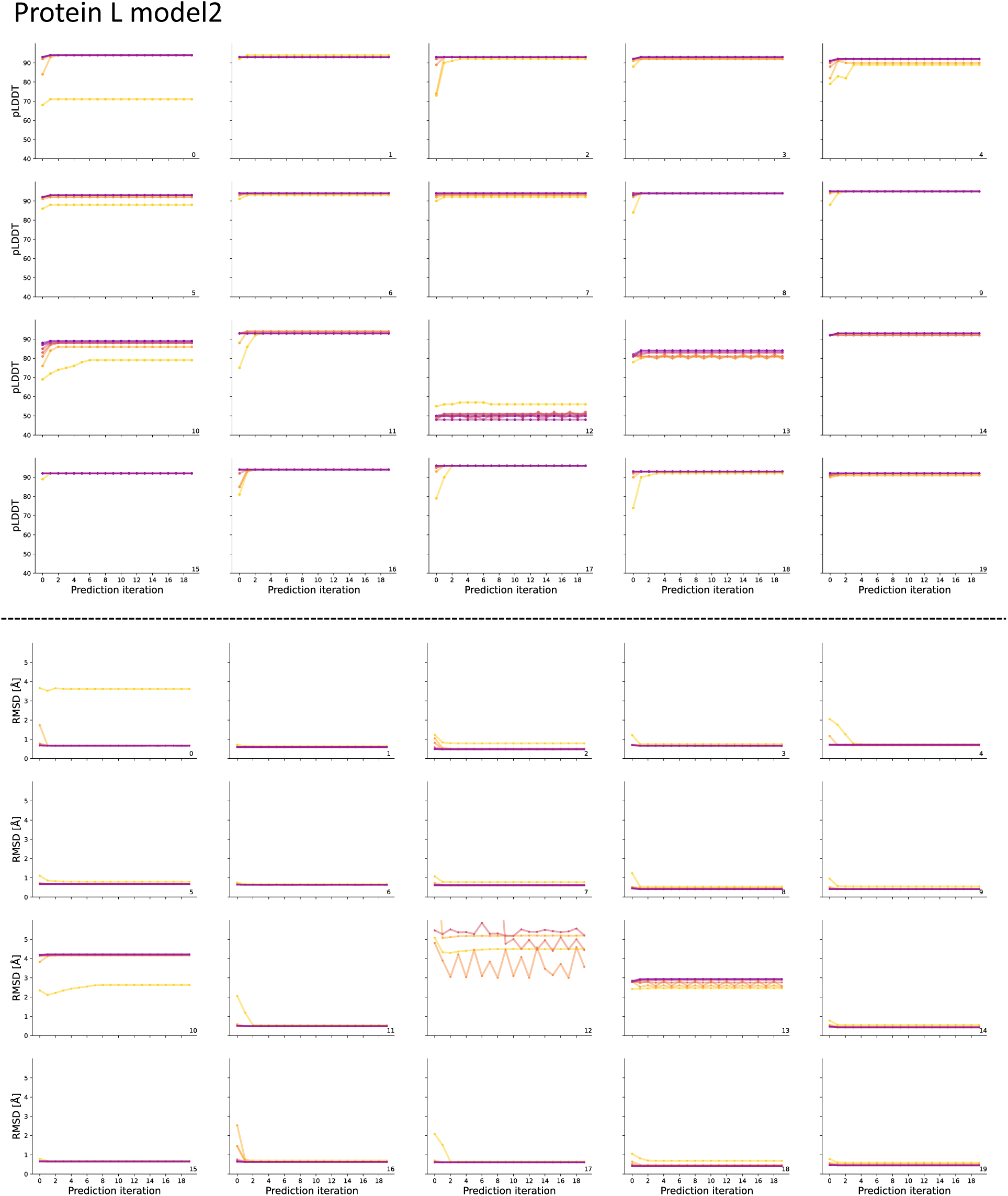
Iterative predictions for 20 ProteinMPNN designed sequences of protein L measured by pLDDT (top) and RMSD (bottom) from native using model 2 of AF2.

**Figure 14: Supplementary figure 8.**
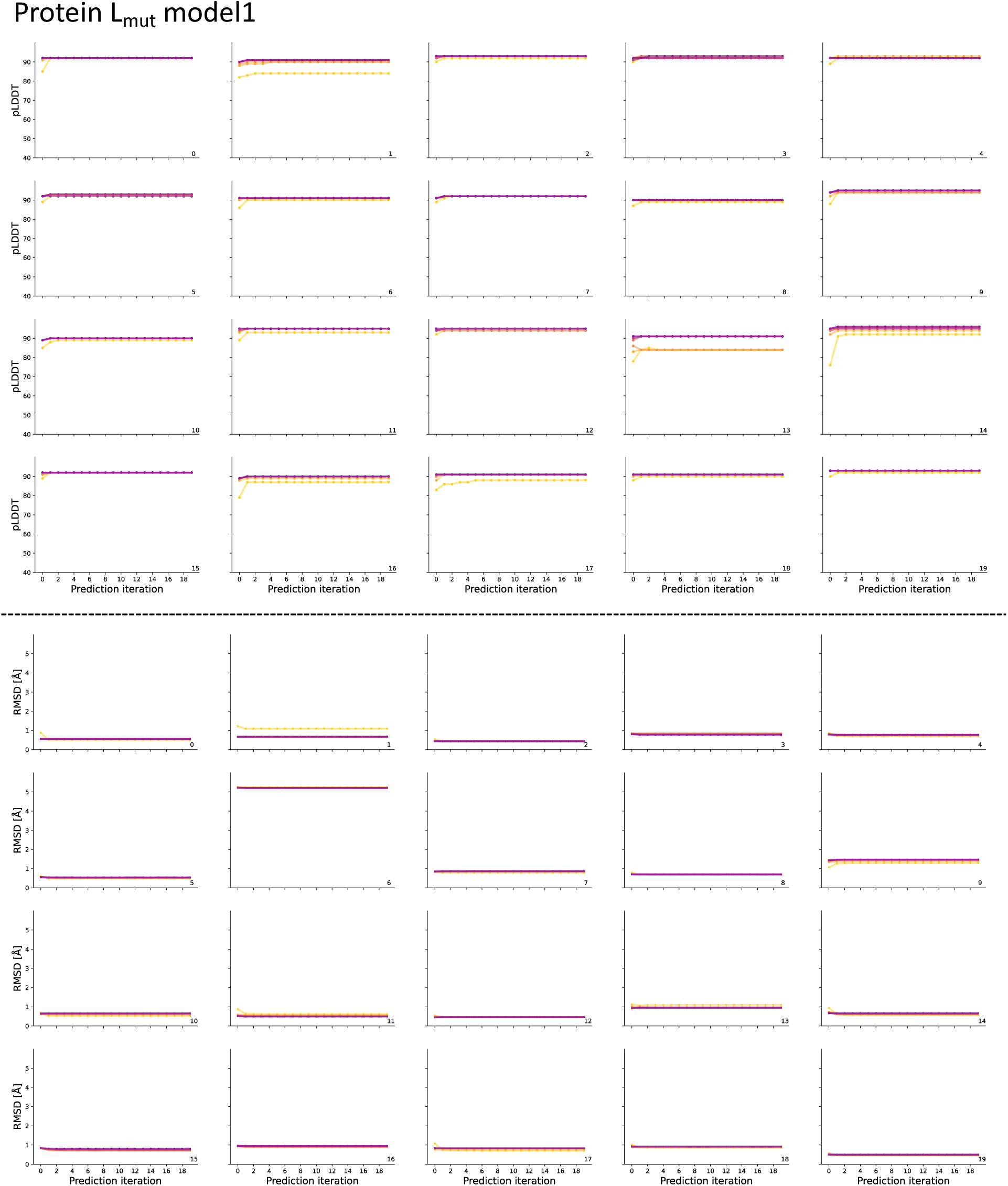
Iterative predictions for 20 ProteinMPNN designed sequences of protein L mutant measured by pLDDT (top) and RMSD (bottom) from native using model 1 of AF2.

**Figure 15: Supplementary figure 9.**
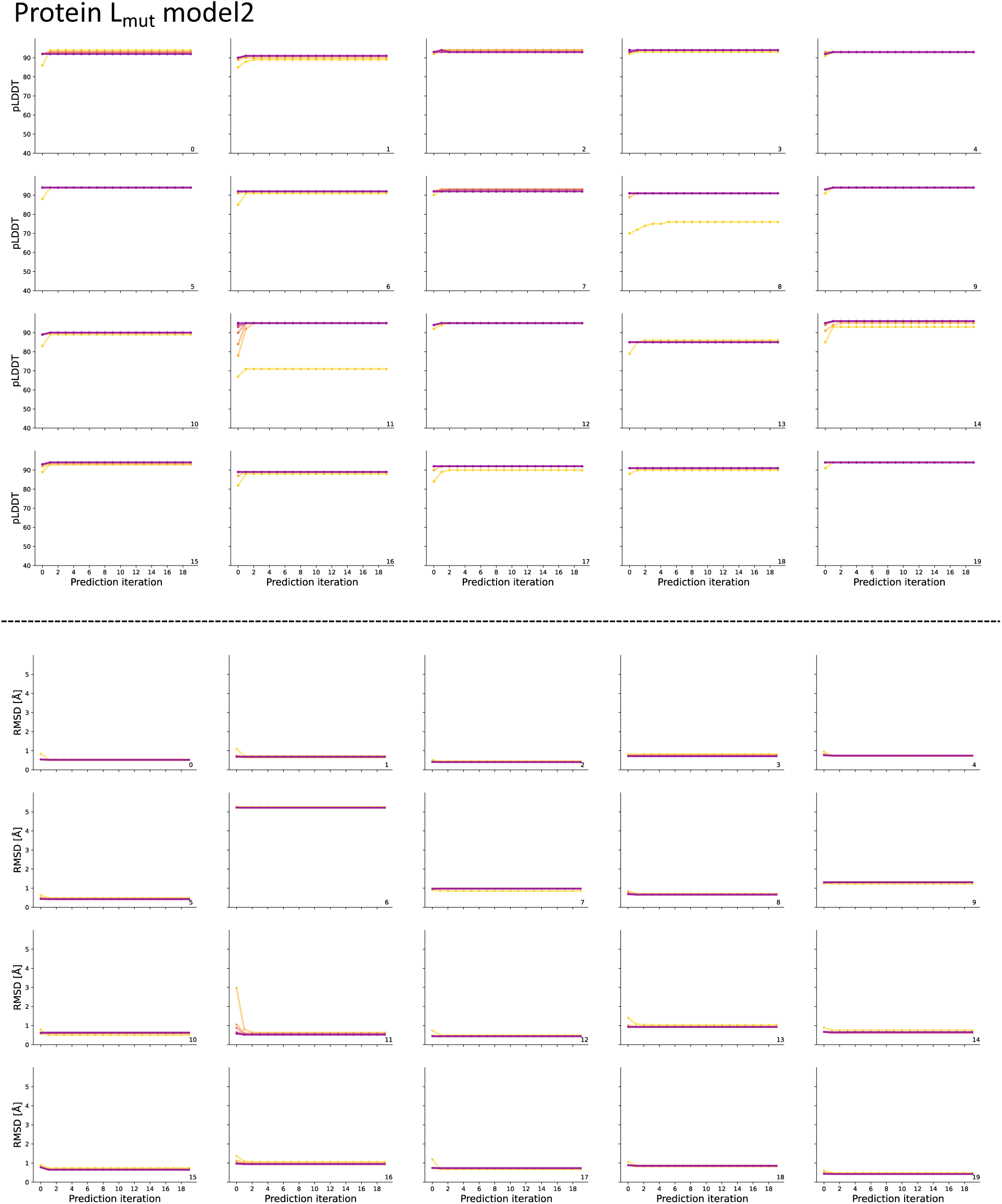
Iterative predictions for 20 ProteinMPNN designed sequences of protein L mutant measured by pLDDT (top) and RMSD (bottom) from native using model 2 of AF2.

**Figure 16: Supplementary figure 10.**
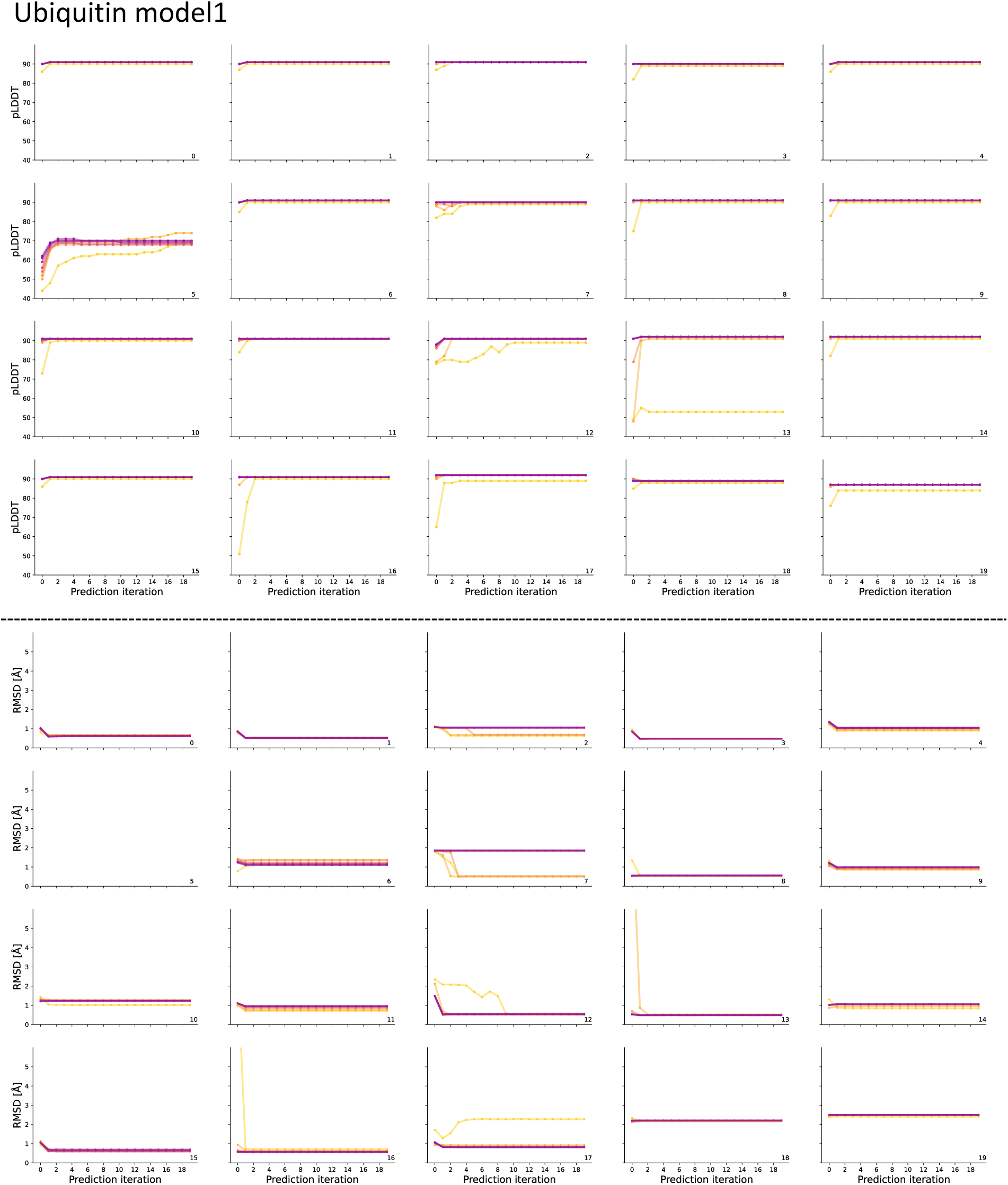
Iterative predictions for 20 ProteinMPNN designed sequences of ubiquitin measured by pLDDT (top) and RMSD (bottom) from native using model 1 of AF2.

**Figure 17: Supplementary figure 11.**
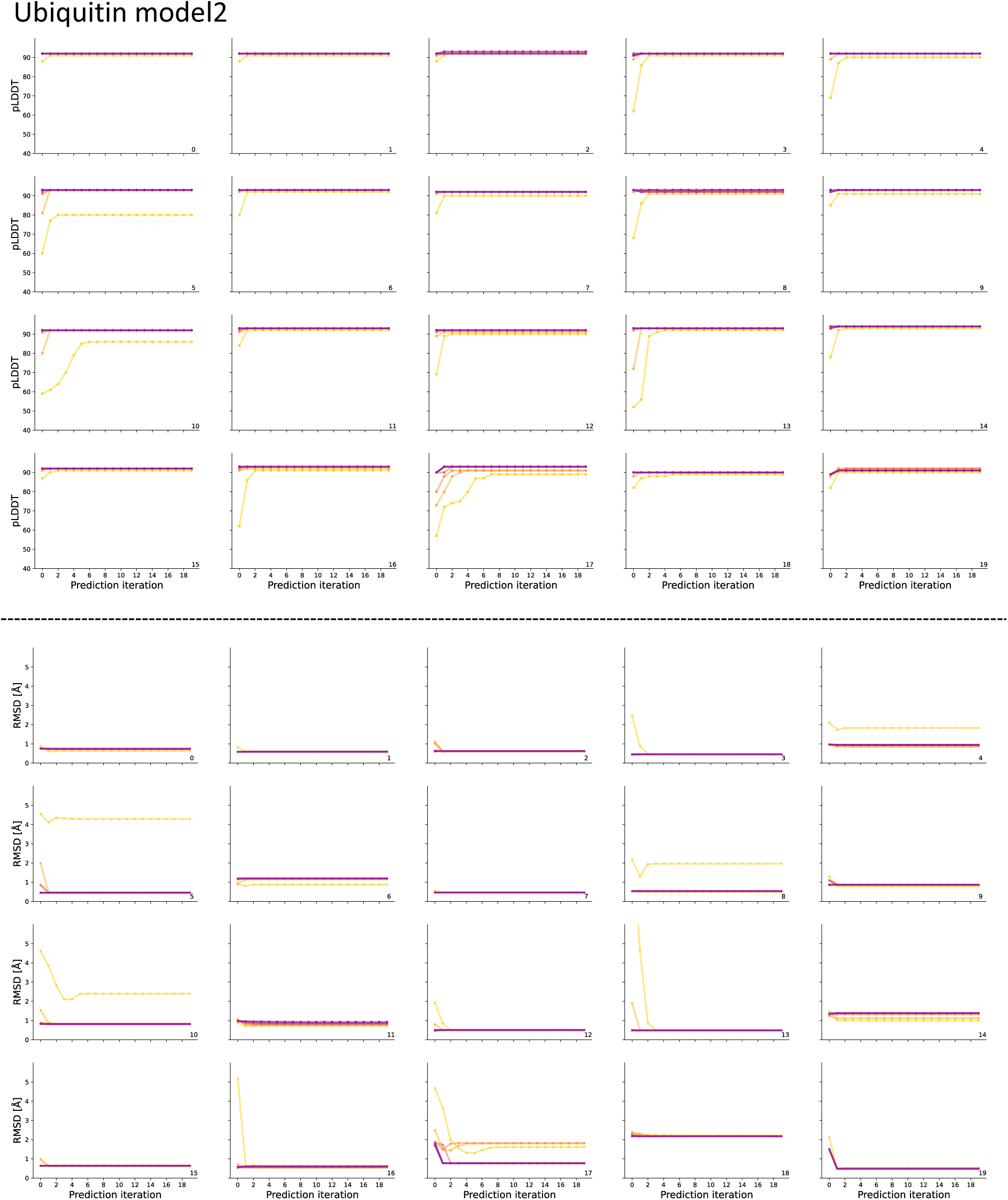
Iterative predictions for 20 ProteinMPNN designed sequences of ubiquitin measured by pLDDT (top) and RMSD (bottom) from native using model 2 of AF2.

**Figure 18: Supplementary figure 12.**
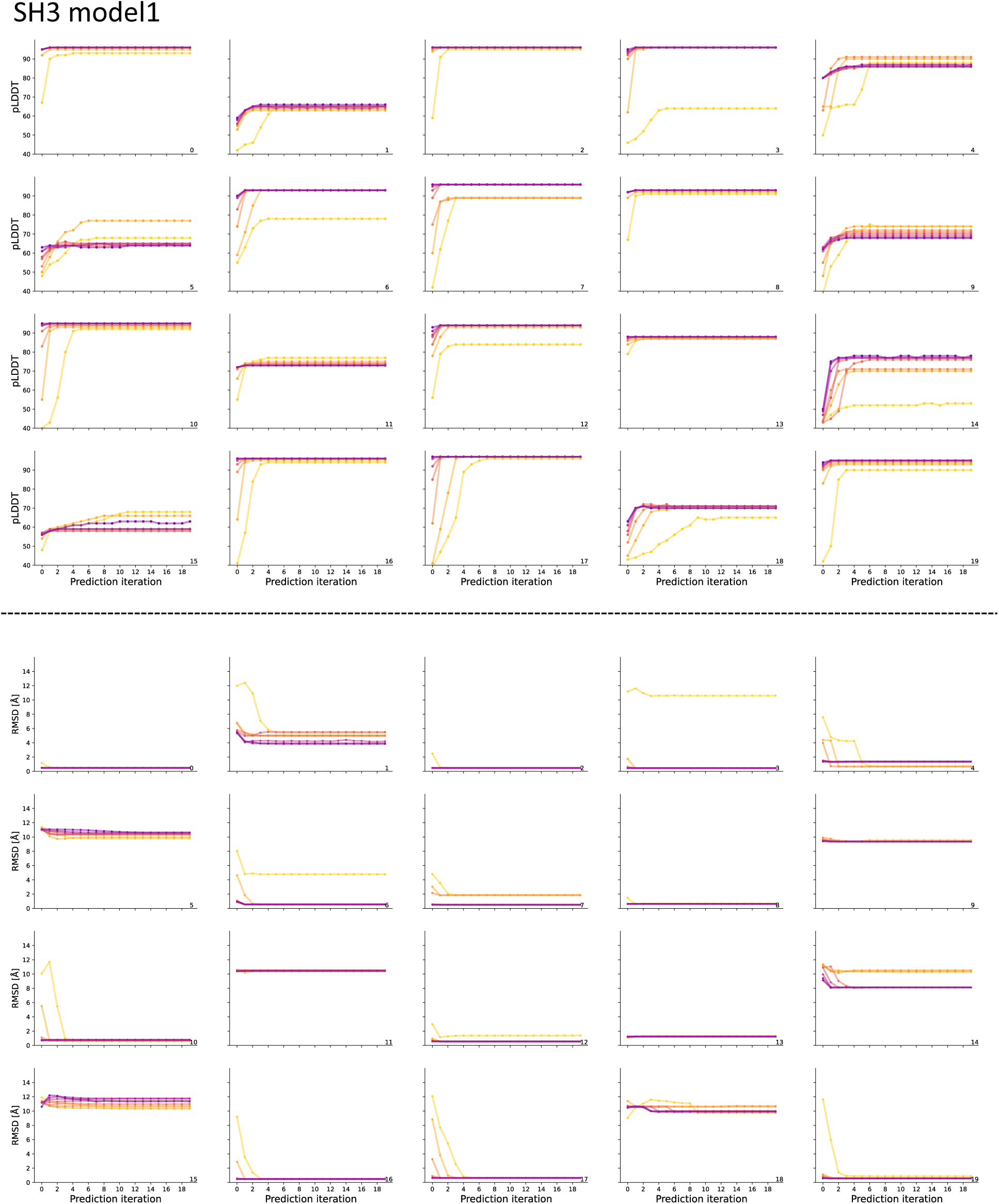
Iterative predictions for 20 ProteinMPNN designed sequences of SH3 measured by pLDDT (top) and RMSD (bottom) from native using model 1 of AF2.

**Figure 19: Supplementary figure 13.**
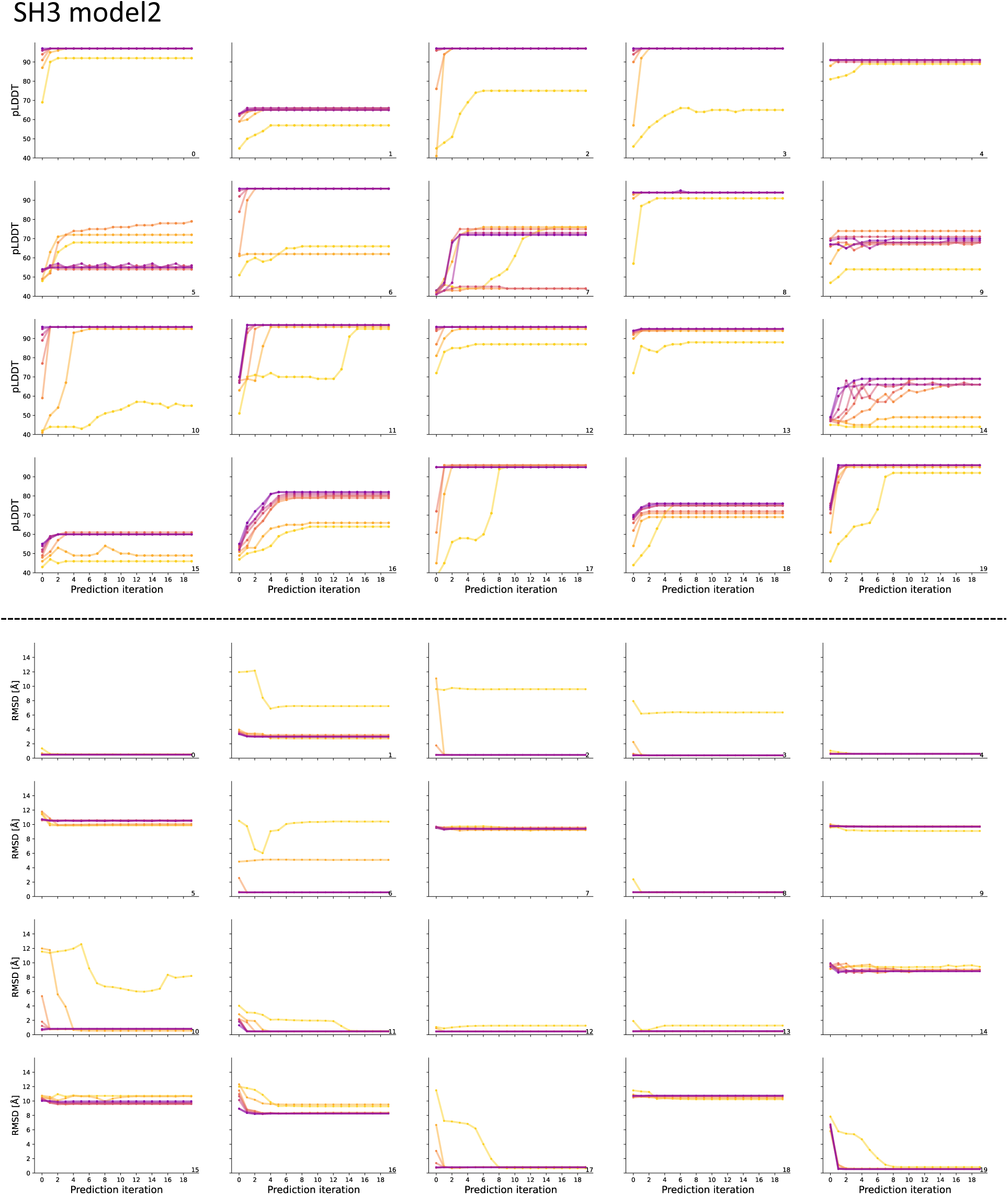
Iterative predictions for 20 ProteinMPNN designed sequences of SH3 measured by pLDDT (top) and RMSD (bottom) from native using model 2 of AF2.

**Figure 20: Supplementary figure 14.**
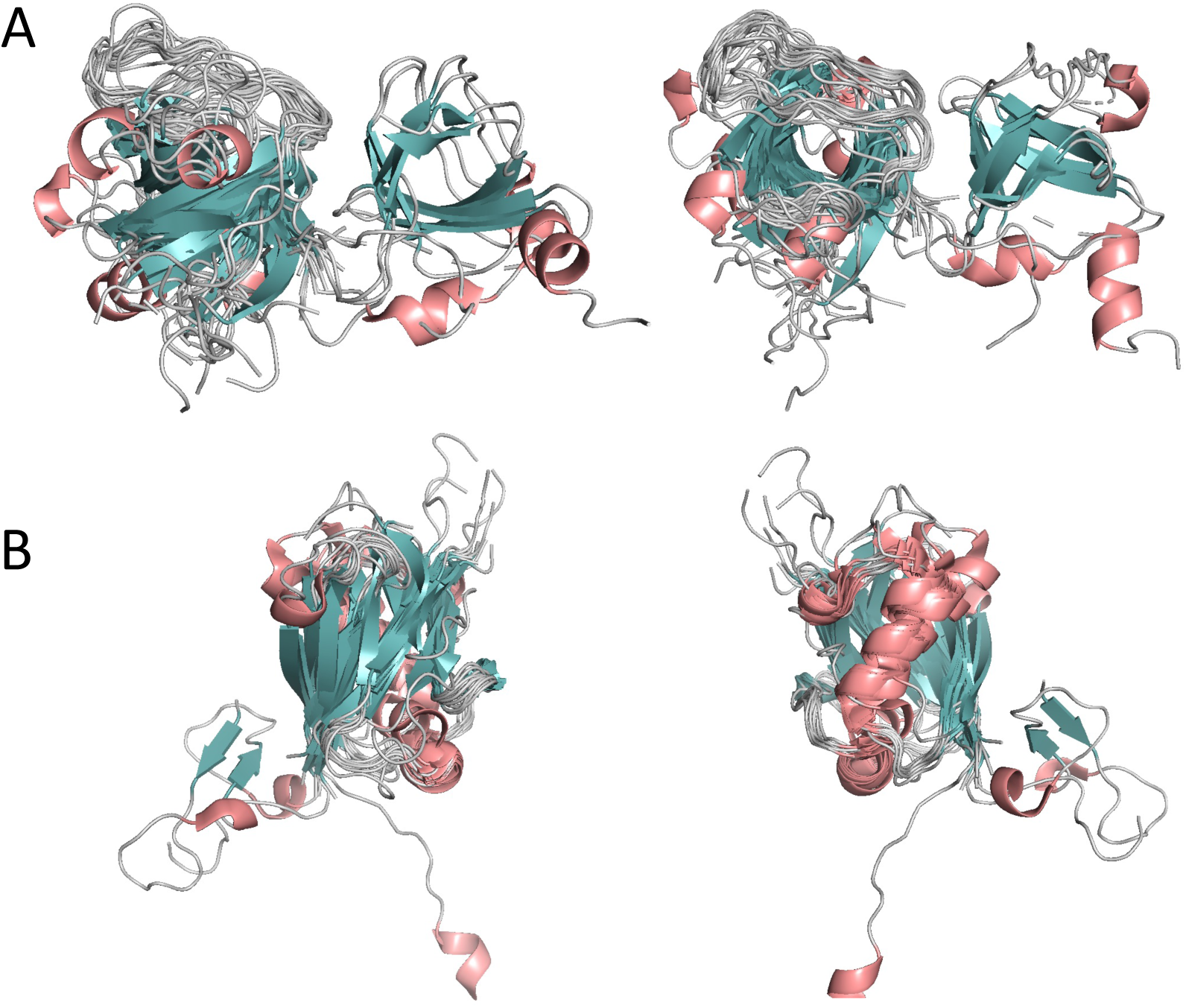
A. Structural alignments of SH3 like proteins (left is front view and right is top view); B. Structural alignments of ubiquitin like proteins (left is front view and right is back view).

**Figure 21: Supplementary figure 15.**
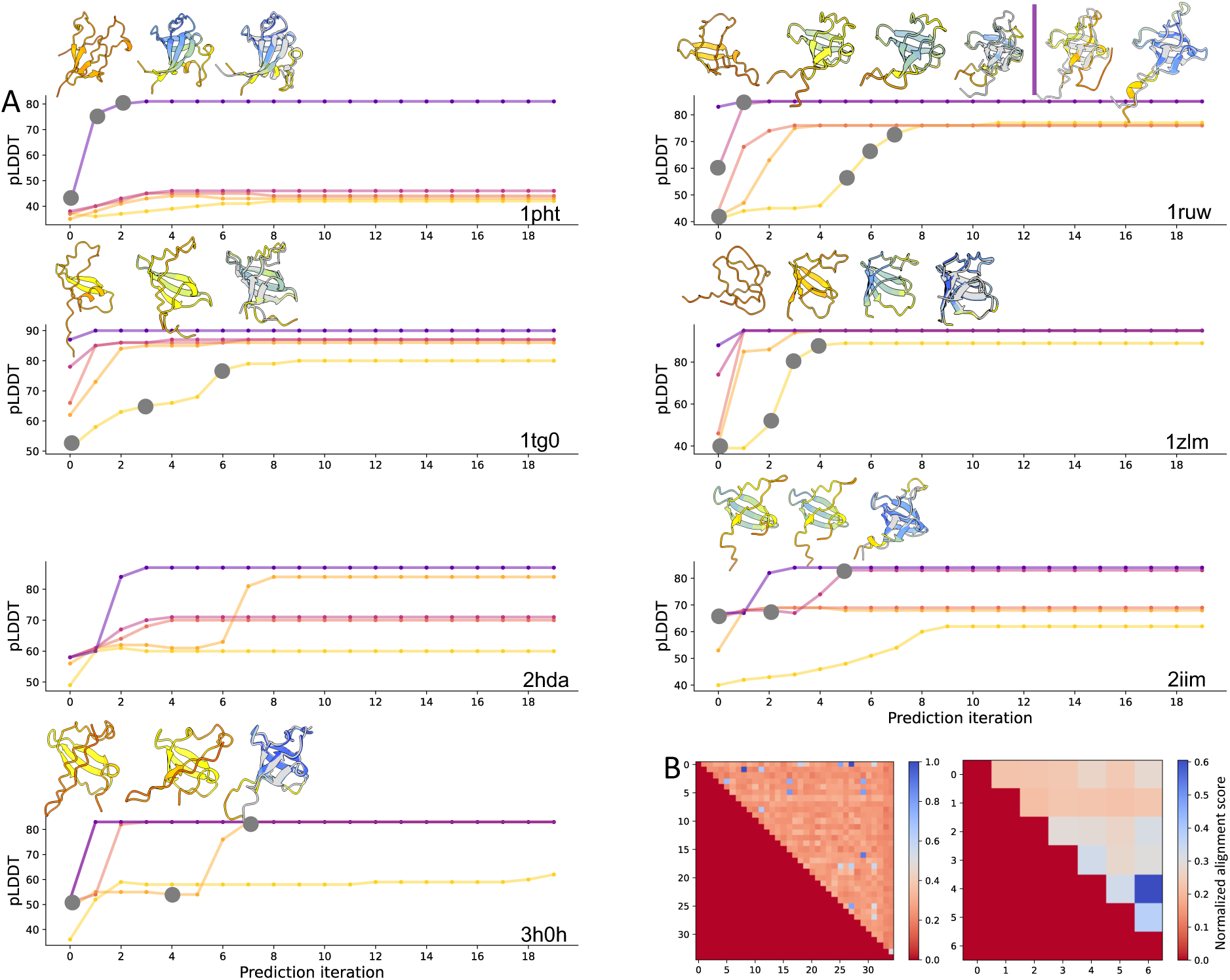
A. Iterative structure predictions of SH3 like proteins which fold into structure less than 3 Å from native state (colored in grey). Representative structures are shown for the iteration successfully finds native structure with the least recycling. B. Pairwise sequence similarity for all SH3 like protein sequences (left) and sequences that find native structures (right).

**Figure 22: Supplementary figure 16.**
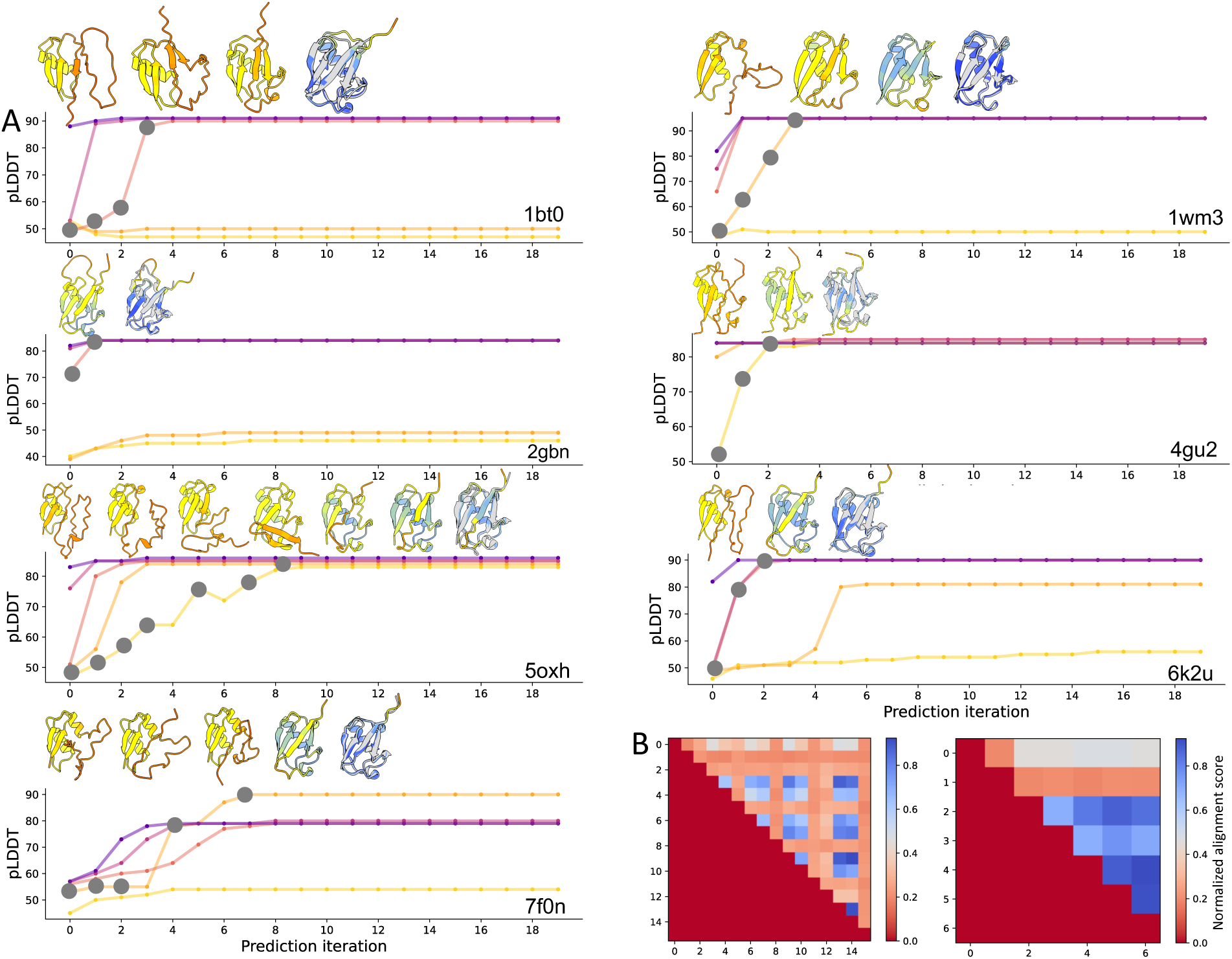
A. Iterative structure predictions of ubiquitin like proteins which fold into structure less than 3 Å from native state (colored in grey). Representative structures are shown for the iteration successfully finds native structure with the least recycling. B. Pairwise sequence similarity for all ubiquitin like protein sequences (left) and sequences that find native structures (right).

